# Caveolae mediated endocytosis of VLDL particles in macrophages requires NPC1 and STARD3 for further lysosomal processing

**DOI:** 10.1101/2021.12.16.473074

**Authors:** Lei Deng, Frank Vrieling, Rinke Stienstra, Guido Hooiveld, Anouk L. Feitsma, Sander Kersten

**Affiliations:** Nutrition, Metabolism and Genomics Group, Division of Human Nutrition and Health, Wageningen University, Stippeneng 4, 6708 WE Wageningen, the Netherlands; FrieslandCampina, Stationsplein 4, 3818 LE Amersfoort, the Netherlands; Department of Internal Medicine, RadboudUMC, Nijmegen, The Netherlands

**Keywords:** Very low-density lipoproteins, endocytosis, caveolae, LPL, Macrophages

## Abstract

Macrophages accumulate triglycerides under certain pathological conditions such as atherosclerosis. Triglycerides are carried in the bloodstream as part of very low-density lipoproteins (VLDL) and chylomicrons. How macrophages take up and process VLDL-lipids is not very well known. Here, using VLDL-sized triglyceride-rich emulsion particles, we aimed to study the mechanism by which VLDL-triglycerides are taken up, processed, and stored in macrophages. Our results show that macrophage uptake of emulsion particles mimicking VLDL (VLDLm) is dependent on lipoproteins lipase (LPL) and requires the lipoprotein-binding C-terminal domain of LPL but not the catalytic N-terminal domain. Subsequent internalization of VLDLm-triglycerides by macrophages is carried out by caveolae-mediated endocytosis, followed by triglyceride hydrolysis catalyzed by lysosomal acid lipase. Transfer of lysosomal fatty acids to the ER for subsequent storage as triglycerides is mediated by Stard3, whereas NPC1 was found to promote the extracellular efflux of fatty acids from lysosomes. Our data provide novel insights into how macrophages process VLDL-derived triglycerides and suggest that macrophages have the remarkable capacity to excrete part of the internalized triglycerides as fatty acids.

**Summary:** Triglyceride-rich lipoproteins and their remnants contribute to atherosclerosis, possibly by carrying remnant cholesterol and/or by exerting a pro-inflammatory effect on macrophage. Nevertheless, little is known about how macrophages process triglyceride-rich lipoproteins. We show that uptake by macrophages of VLDL-like particles is dependent on the enzyme lipoproteins lipase via its C-terminal domain. Subsequent internalization of VLDL-triglycerides by macrophages is carried out by caveolae-mediated endocytosis, followed by hydrolysis by lysosomal acid lipase. Transfer of lysosomal fatty acids to the ER for lipid storage is mediated by Stard3, while NPC1 promotes the extracellular efflux of fatty acids. Our data provide novel insights into how macrophages process VLDL-derived triglycerides and suggest that macrophages have the remarkable capacity to excrete internalized triglycerides as fatty acids.

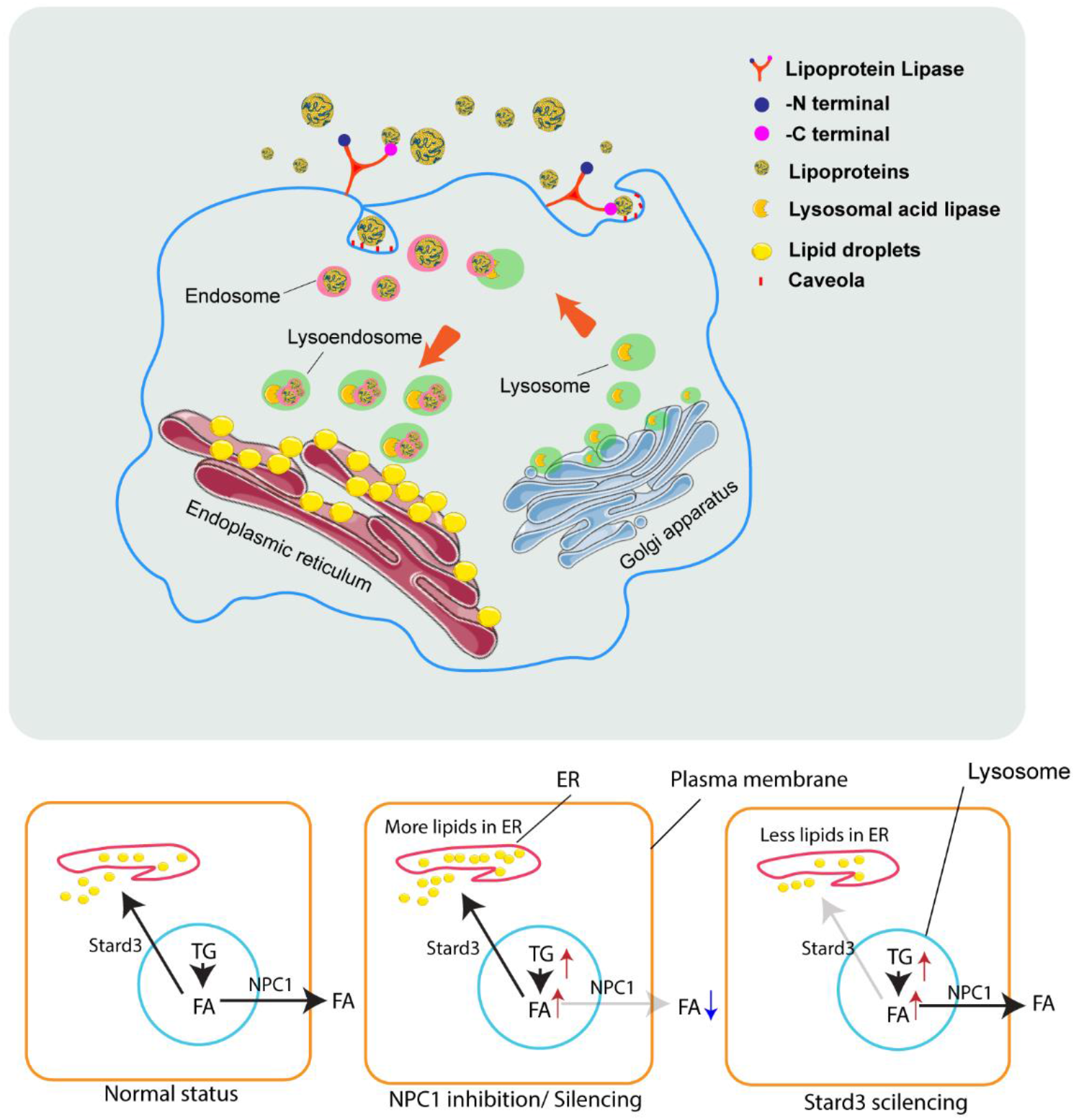

## Introduction

Lipids are essential for all cells, either as structural component, signaling molecule, or fuel source. Lipids are transported through the bloodstream as part of lipoproteins. Whereas cholesterol is mainly carried in high-density and low-density lipoproteins, triglycerides are predominantly transported by chylomicrons and very low-density lipoproteins (VLDLs). Chylomicrons have a diameter of 75-600 nm, carry dietary triglycerides, and are produced by enterocytes [1]. Conversely, VLDLs have an average diameter of 30-80 nm, carry triglycerides that are produced endogenously, and are synthesized in the liver [2]. In the fasted state, triglycerides are present in the blood almost entirely as part of VLDL and its remnant lipoproteins.

Macrophages are innate immune cells that form the frontline in the host defense against pathogens. They are specialized in the detection, phagocytosis, and destruction of bacteria and other harmful organisms. In addition, macrophages can present antigens and regulate inflammation by releasing cytokines. Besides phagocytizing and neutralizing pathogens, macrophages can also scavenge lipids, which after uptake can be stored in specialized organelles called lipid droplets. How macrophages scavenge VLDL and chylomicrons, and how the associated lipids are internalized and processed has not been well characterized.

Lipid uptake and storage in macrophages has primarily been investigated in the context of atherosclerosis [3]. Macrophages take up oxidized LDL, which is considered a key event in the pathogenesis of atherosclerotic lesions [4]. In the past decades, evidence has been accumulating that apart from LDL particles, triglyceride-rich lipoproteins and their remnants also contribute to atherosclerosis, possibly by carrying remnant cholesterol and/or by exerting a pro-inflammatory effect on macrophage [5]. However, in contrast to uptake of oxidized LDL, little is known about how macrophages process these triglyceride-rich particles. Consistent with their purported roles in atherosclerosis, VLDL and its remnants can be taken up and retained in the intima, where they can interact with macrophages [6].

The uptake of oxidized LDL and cholesterol by macrophages is mediated by a group of structurally unrelated molecular pattern recognition receptors referred to as scavenger receptors, including SR-B1, CD36, SR-AI and SR-AII [7]. By comparison, little is known about the uptake mechanism of VLDL-derived triglycerides (VLDL-TG) by macrophages. It has been shown that the uptake of VLDL-TG in cultured macrophages is promoted by LPL [8]. Besides via its lipolytic function, LPL may enhance lipid uptake by functioning as a molecular bridge between VLDL and lipoprotein receptors and/or heparan sulfate proteoglycans [9][10]. Other steps in the uptake of VLDL-TG by macrophages remain poorly defined.

Endocytosis describes the transport of extracellular substances or particles into cells and is tightly related to the biological function of macrophages. The endocytosis pathway can be classified into several types: clathrin-dependent endocytosis, clathrin-independent endocytosis, pinocytosis, and phagocytosis [11]. The importance of clathrin-mediated endocytosis in cellular uptake of lipids is well established. Indeed, after binding to the LDL receptor, cellular uptake of LDL is mediated by binding of LDL to the LDL receptor, followed by formation of clathrin-coated pits and subsequent delivery of the lipid cargo to the lysosomes [12]. Receptor-mediated endocytosis also mediates uptake of native or modified LDL and remnant lipoproteins in macrophages as key step in the pathogenesis of atherosclerosis [13]. In hepatocytes, receptor-mediated endocytosis is required for the uptake of VLDL- and chylomicron remnants [14]. However, little is known about the potential involvement of endocytosis in the uptake of VLDL-TG by macrophages.

After uptake by cells, LDL is degraded in lysosomes, releasing free cholesterol. The cholesterol binds to Niemann-Pick type C 2 (NPC2) before being shuttled out of the lysosome via NPC1 [15] [16][17]. Part of the cholesterol transported via this pathway may go to the plasma membrane, while another portion may go to the endoplasmic reticulum (ER). The lysosomal membrane protein Stard3 transports the cholesterol directly from the lysosome to the ER by interacting with vesicle-associated protein-associated protein (VAP)A/B on the ER membrane [18]. Interestingly, Stard3 also promotes the reverse transport of cholesterol from the ER to the lysosome [19], as well as the transfer of cholesterol from lysosome to mitochondria [20]. While there is thus substantial insight into how cholesterol is processed, transported, and stored in macrophages, our mechanistic understanding of how VLDL-TG are processed in macrophages is very limited.

Here, using VLDL-sized triglyceride-rich emulsion particles, we aimed to study the mechanism by which VLDL-TG are taken up and processed in macrophages.

## Results

### VLDL-sized lipid emulsion particles are taken up by macrophages

Using a microfluidizer, triglyceride-rich emulsion particles were prepared with a mean diameter of 60 nm, referred to as VLDL-mimic (VLDLm) (Figure 1a)[21]. Treatment of human primary macrophages with these particles for 6 hours led to marked lipid accumulation, as visualized using BODIPY 493/503 staining (Figure 1b) and quantified using flow cytometry (Figure 1c and Figure S1b). Treatment with VLDLm also significantly increased expression of lipid-sensitive genes as measured by Realtime-qPCR (Figure 1d and Figure S1c). Heatmaps (Figure 1e and Figure S1d) and a volcano plot (Figure 1f) using all genes with q value less than 0.05 for any comparison using RNA sequencing clearly showed the marked effect of the lipid emulsion particles on gene expression in macrophages. Interestingly, VLDLm reduced the expression of cholesterol biosynthesis genes (GO:0006695), which is consistent with the suppressive effect of unsaturated fatty acids on genes involved in cholesterol synthesis [22][23], and further illustrates the significant impact of the lipid emulsion particles on lipid metabolism in macrophages (Figure 1g). In agreement with previous studies [24][25][26][27], VLDLm also modulated genes involved in the ER stress response (GO:0034976) and the inflammatory response (GO:0006954) (Figure S1g-h).

**Figure 1.**
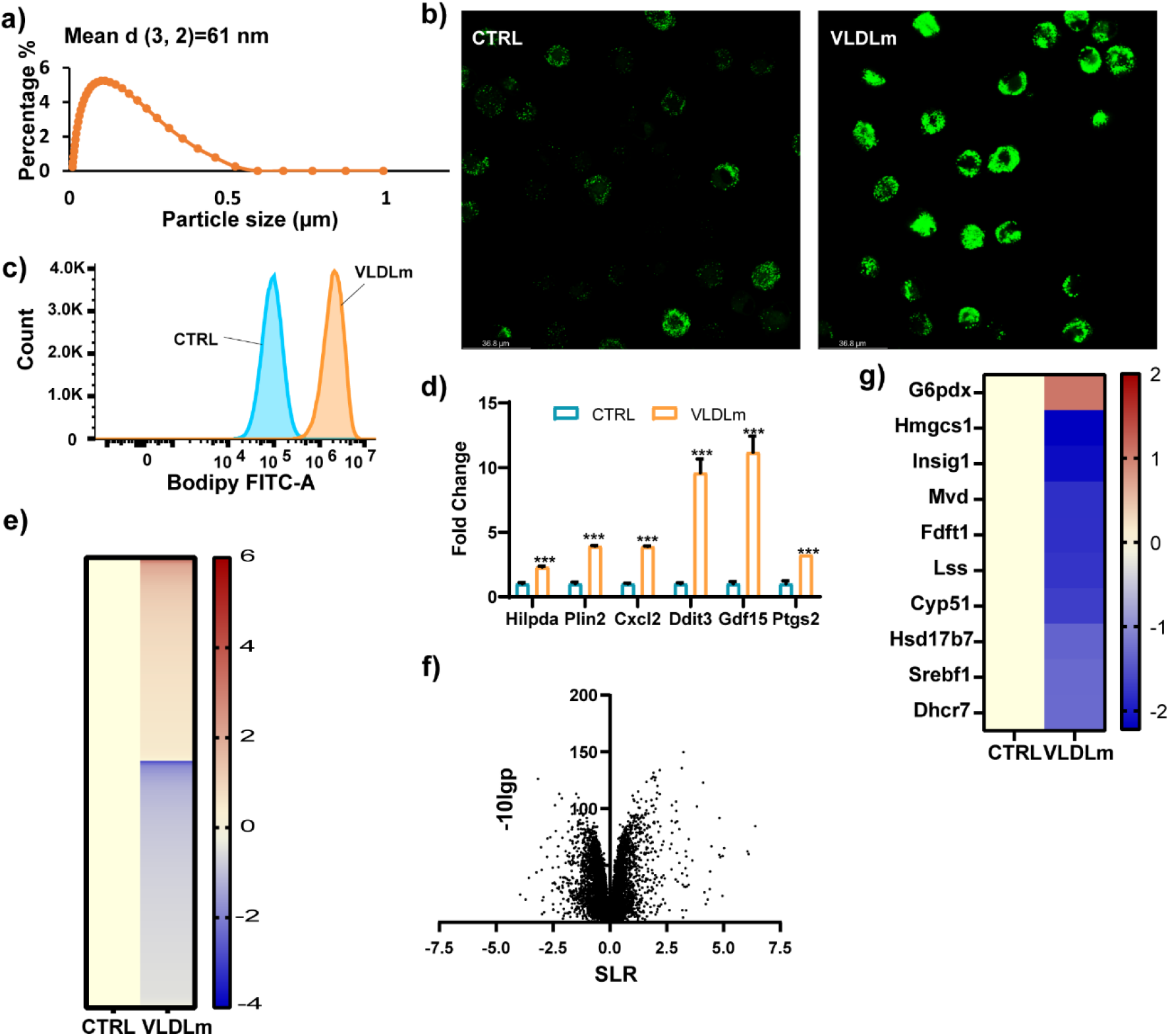
VLDLm promotes lipid accumulation in cultured macrophages. a) The particle size distribution of VLDLm as determined by mastersizer 3000. b) BODIPY 493/503 staining of intracellular neutral lipids in human primary macrophages treated with 0.5 mM VLDLm for 6 hours. c) Mean fluoresce intensity (FITC-A) measured by flow cytometry of mouse RAW 264.7 macrophages treated with 1 mM VLDLm for 6 hours. d) mRNA expression of lipotoxic marker genes in RAW 264.7 macrophages treated with 1mM VLDLm for 6 hours. e) Heatmap and f) Volcano plot of RNAseq data of RAW 264.7 macrophages treated with 1 mM VLDLm for 6 hours. g) Heatmap plotted with differentially expressed genes (p<0.01, SLR>1) involved in the cholesterol synthesis pathway. Bar graphs were plotted as mean±SD. Significance was analysed using Student’s t-test; *P < 0.05, **P<0.01, ***P<0.001, ***P<0.0001.

### LPL is required for uptake of VLDLm-TG by macrophages

To determine whether LPL is required for uptake of VLDLm-TG by macrophages, we treated RAW 264.7 macrophages with heparin, which releases LPL from the cell surface. As expected, treatment with heparin markedly reduced macrophage LPL content (Figure 2a). The LPL remaining after heparin treatment likely represents the functionally inactive intracellular LPL pool. Consistent with a role of LPL in uptake of VLDLm-TG by macrophages, heparin treatment reduced lipid accumulation in VLDLm-treated macrophages (Figure 2b-c).

**Figure 2.**
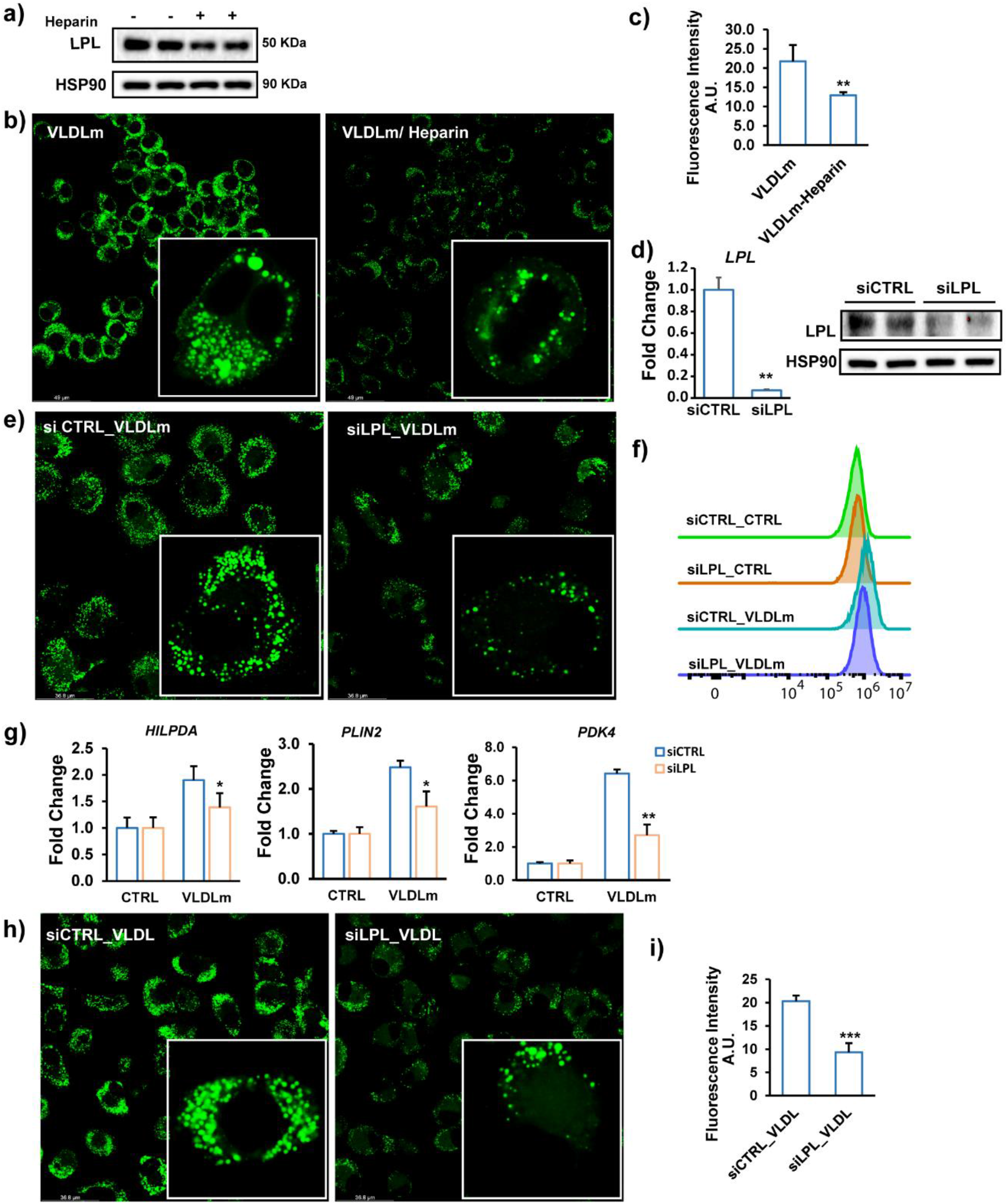
LPL mediates VLDL uptake in cultured macrophages. a) LPL protein levels in RAW 264.7 macrophages treated with 50 UI/ml Heparin for 2 hours. b) BODIPY 493/503 staining of intracellular neutral lipids in RAW 264.7 macrophages treated with 1mM VLDLm for 6 hours in the presence or absence of 50 UI/ ml human heparin. c) Quantification of the fluorescence images by ImageJ. d) LPL mRNA and protein levels in human primary macrophages treated with control siRNA or LPL siRNA for 48 hours. e) BODIPY 493/503 staining of intracellular neutral lipids in human macrophages treated with siCTRL or siLPL for 48 hours followed by treatment with 0.5 mM VLDLm for 6 hours. f) Mean fluorescence intensity quantified by flow cytometry. g) mRNA expression of selected lipid-sensitive genes. h) BODIPY 493/503 staining of intracellular neutral lipids in human macrophages treated with siCTRL or siLPL for 48 hours followed by treatment with 0.5 mM human plasma isolated VLDL for 6 hours. i) Mean fluorescence intensity quantified by flow cytometry. The bar graphs were plotted as mean±SD. Asterisk indicates significantly different from control according to Student’s t-test. *p<0.05, **p<0.01, ***p<0.001.

To further assess the role of LPL in macrophage uptake of VLDLm, we silenced LPL in human primary macrophages using siRNA (Figure 2d). Silencing of LPL markedly reduced cellular lipid accumulation in VLDLm-treated macrophages, as visualized by BODIPY 493/503 staining (Figure 2e) and supported by flow cytometric analysis (Figure 2f). Consistent with lower lipid uptake, the induction of lipid sensitive genes *Hilpda, Plin2* and *Pdk4* by VLDLm was significantly reduced by LPL silencing (Figure 2g). A similar suppressive effect of LPL silencing on lipid accumulation was observed in macrophages treated with human VLDL (Figure 2h-i). Overall, these data indicate that LPL is necessary for macrophage uptake of VLDLm-TG.

### Uptake of VLDLm-TG by macrophages is dependent on the C-terminal portion of LPL

Based on our and other previous data [24][28][29], we hypothesized that the catalytic function of LPL is essential for macrophages uptake of VLDLm-TG. However, no significant changes were observed in lipid accumulation in VLDLm-treated RAW 264.7 cells after the application of GSK264220A, a catalytic inhibitor for LPL [23] (Figure 3a-b). Also, addition of an antibody directed against the catalytic N-terminal portion of human LPL did not noticeably influence lipid accumulation in human monocytes-derived macrophages (Figure 3c-d). These data suggest that uptake of VLDLm-TG by macrophages does not require the catalytic function of LPL.

**Figure 3.**
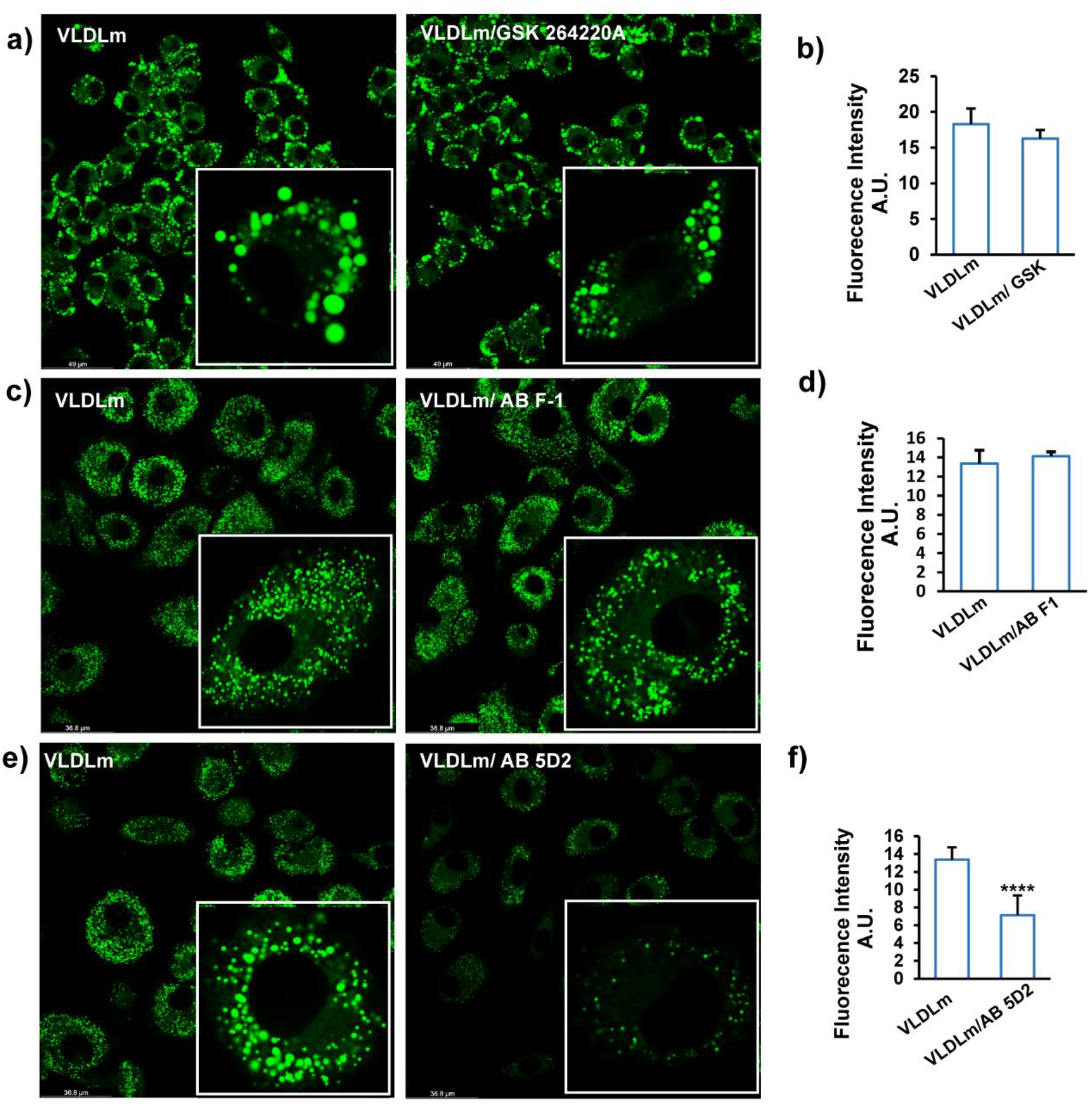
The c-terminal portion of LPL mediates VLDLm uptake in cultured macrophages. a) BODIPY 493/503 staining of RAW 264.7 macrophages treated with 1mM VLDLm for 6 hours in the presence or absence of 0.2 μM of the catalytic LPL inhibitor GSK264220. b) Mean fluorescence intensity quantified by flow cytometry. c) BODIPY 493/503 staining of RAW 264.7 macrophages treated with 1mM VLDLm for 6 hours in the presence or absence of antibody F1 targeting the N-terminal portion of LPL (2 μg/ml). d) Mean fluorescence intensity quantified by flow cytometry. e) BODIPY 493/503 staining of RAW 264.7 macrophages treated with 1mM VLDLm for 6 hours in the presence or absence of antibody 5D2 targeting the C-terminal of LPL (2 μg/ml), f) Mean fluorescence intensity quantified by flow cytometry. The bar graphs were plotted as mean±SD. Asterisk indicates significantly different from control according to Student’s t-test. ****p<0.0001.

The requirement of macrophage uptake of VLDLm-TG on LPL but not on the catalytic activity of LPL suggest that LPL may participate in the binding of the VLDLm particles to the macrophage surface. To verify this notion, human primary macrophages were co-treated with VLDLm and an antibody (5D2) directed against the C-terminal portion of LPL, which mediates the binding of TG-rich lipoproteins [25]. Interestingly, the C-terminal hLPL antibody marked decreased intracellular lipid accumulation, supporting the notion that the C-terminal region of LPL is required for macrophage uptake of VLDLm-TG and suggesting that LPL’s role in macrophage uptake of VLDLm-TG is more as a receptor than as an enzyme (Figure 3e-f).

### Macrophage uptake of VLDLm-TG is mediated by caveola-mediated endocytosis

Next, we examined whether uptake of VLDLm-TG was mediated by endocytosis. In agreement with this notion, early endosomes could be observed in RAW264.7 macrophages after loading with VLDLm (Figure 4a). To determine if VLDLm-TG are taken up by macrophages via clathrin- or caveola-mediated endocytosis, RAW264.7 macrophages were treated with VLDLm in conjunction with the endocytosis inhibitors cholopromazine and genistein, which block clathrin- and caveola-mediated endocytosis, respectively [30]. Whereas cholopromazine showed little to no effect, genistein markedly reduced intracellular lipid accumulation (Figure 4b-c), suggesting that VLDLm-TG are taken up via caveola-mediated endocytosis. Genistein similarly reduced intracellular lipid accumulation in human primary macrophages treated with VLDLm (Figure 4f-g).

**Figure 4.**
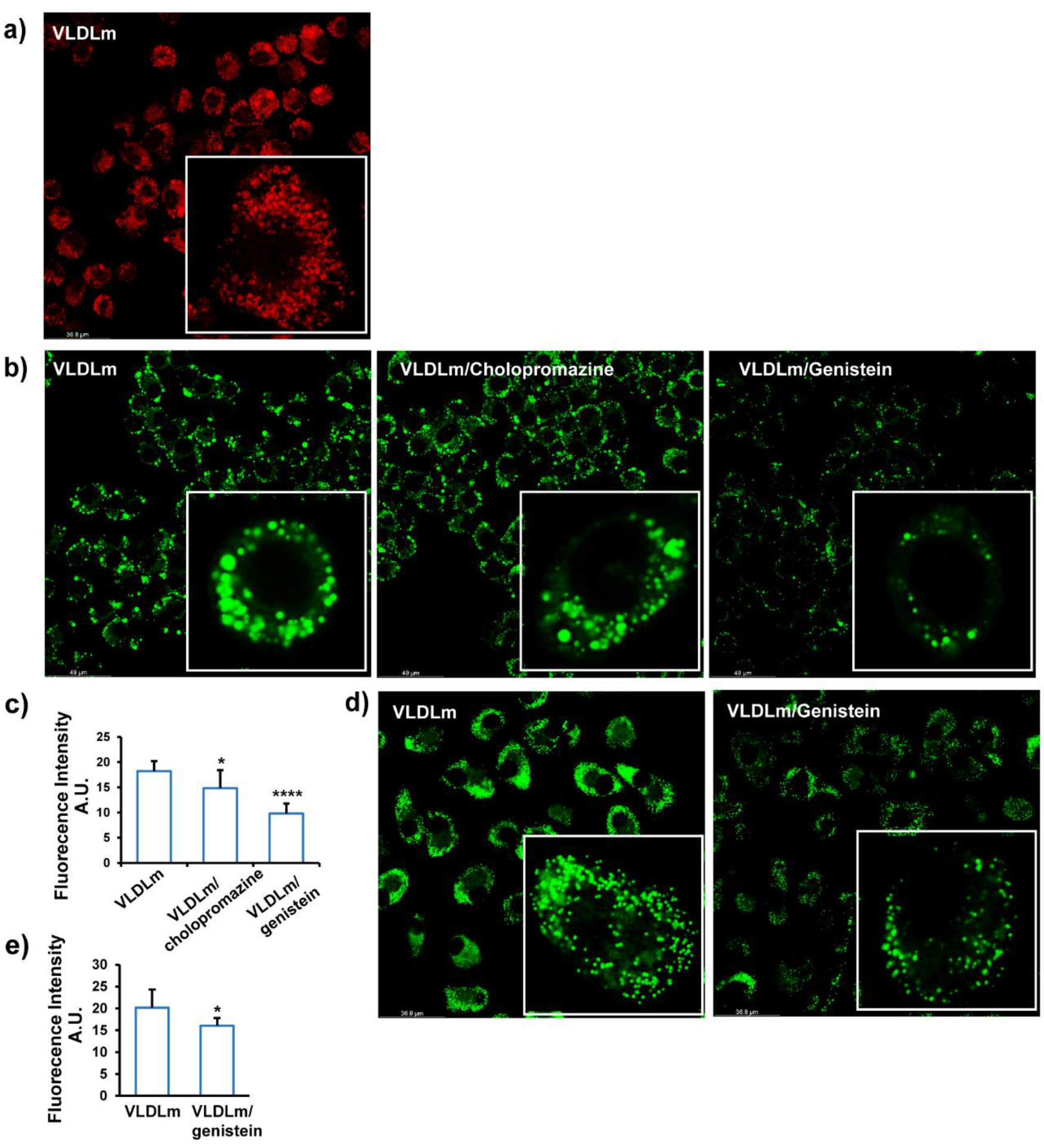
VLDLm are taken up by macrophages via caveola-mediated endocytosis. a) Early endosome staining of RAW 264.7 macrophages treated with 1 mM treatment of VLDLm for 6 hours. b) BODIPY 493/503 staining of RAW 264.7 macrophages treated with 1mM VLDLm for 6 hours in the presence or absence of 10μg/ml chlorpromazine or 200 μM Genistein. c) Mean fluorescence intensity quantified by flow cytometry. d) BODIPY 493/503 staining of human primary macrophages treated with 0.5 mM VLDLm for 6 hours in the presence or absence of 200 μM Genistein. e) Mean fluorescence intensity quantified by flow cytometry. The bar graphs were plotted as mean±SD. Asterisk indicates significantly different from control according to Student’s t-test. *p<0.05.

To further examine the role of caveola in uptake of VLDLm-TG, we silenced the CAV1 and CAV2 genes in human primary macrophages using siRNA (Figure 5a-b). Caveolins, encoded by the CAV1 and CAV2, are the main protein components of caveolae. Silencing of CAV1 and CAV2 markedly decreased intracellular lipid accumulation after loading cells with VLDLm, as visualized by confocal microscopy VLDLm loading (Figure 5c) and quantified by flow cytometry (Figure 5d). A similar marked reduction in intracellular lipid accumulation upon CAV1 (Figure 5e-f) and CAV2 silencing (Figure 5g-h) was observed in human primary macrophages treated with human plasma isolated VLDL.

**Figure 5.**
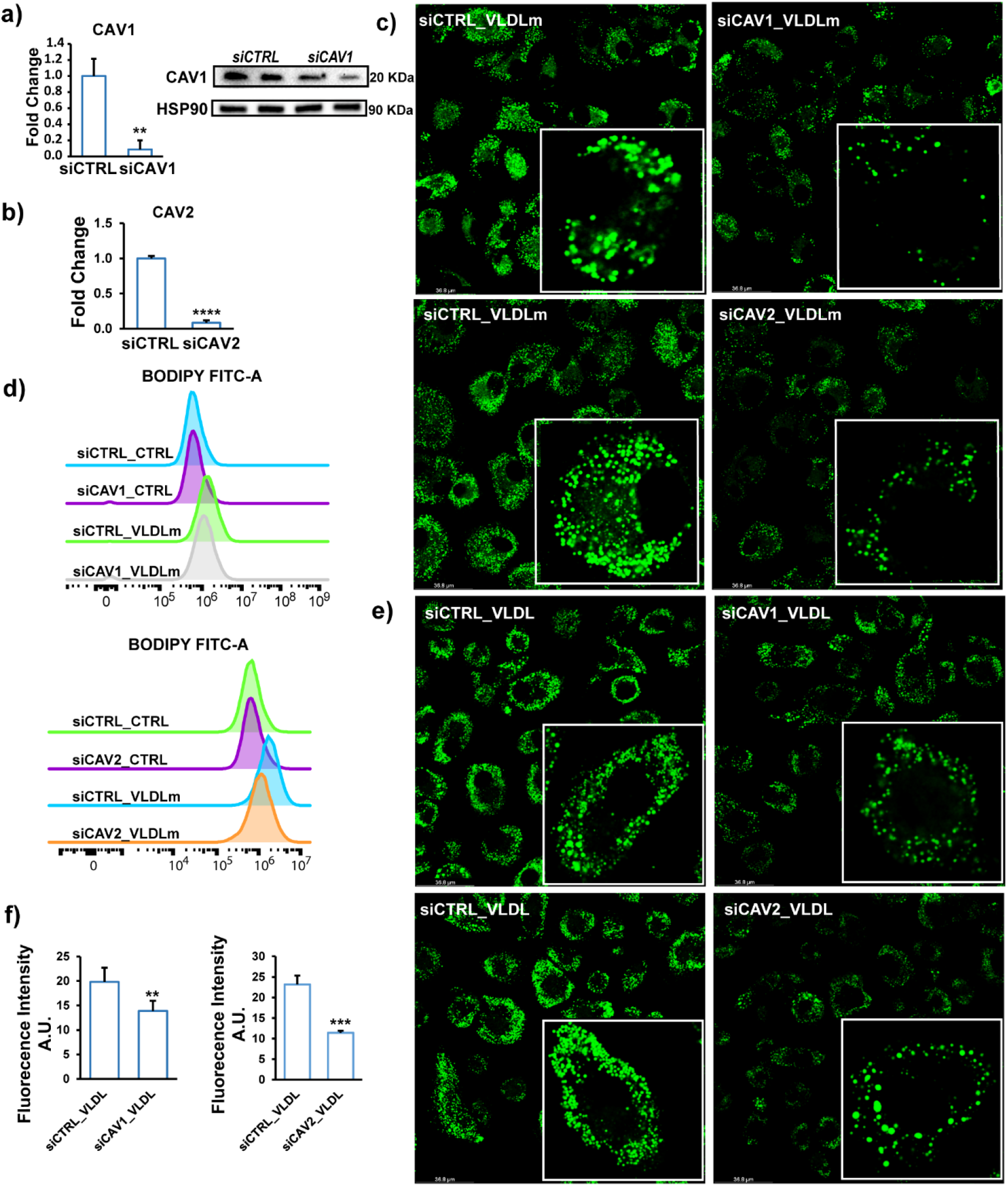
Silencing of Caveolin 1 and 2 impairs uptake of VLDL by macrophages. a) mRNA and protein levels of Caveolin 1 after 48 hour treatment with siCTRL or siCAV1. b) mRNA expression of Caveolin 2 after 48 hour treatment with siCTRL or siCAV2. c) BODIPY 493/503 staining of human macrophages treated with siCTRL, siCAV1, or siCAV2 for 48 hours followed by treatment with 0.5 mM VLDLm for 6 hours. d) Mean fluorescence intensity quantified by flow cytometry. e) BODIPY 493/503 staining of human macrophages treated with siCTRL, siCAV1, or siCAV2 for 48 hours followed by treatment with 0.5 mM human plasma isolated VLDL for 6 hours. f) Mean fluorescence intensity quantified by flow cytometry. The bar graphs were plotted as mean±SD. Asterisk indicates significantly different from control according to Student’s t-test. **p<0.01, ***p<0.001.

Treatment of RAW264.7 macrophages with VLDLm was associated with marked upregulation of genes involved in lysosomal function (Figure 6a). Endocytosed lipids destined for degradation are sorted to lysosomes, where they are digested by the enzyme lysosomal acid lipase. Strikingly, co-treatment of VLDLm-treated human primary macrophages with the LAL inhibitor Lalistat 2 markedly enhanced intracellular lipid accumulation (Figure 6b and Figure 6c). Co-staining of lipids and lysosomes showed that the lipids appeared to be trapped in the lysosomal compartment (Figure 6d). The increase in lipid accumulation by Lalistat 2 was accompanied by reduced expression of the lipid-sensitive genes *HILPDA*, *PDK4*, *PLIN2*, *CXCL2* (Figure 6e). These data suggest that in the presence of LAL inhibitor, endocytosed lipids cannot further be processed and accumulate in lysosomes and also prevents the transcriptional activation of lipid-sensitive genes. Taken together, our results demonstrate that VLDLm-TG are taken up via caveolae-mediated endocytosis and are further degraded via LAL in the lysosome.

**Figure 6.**
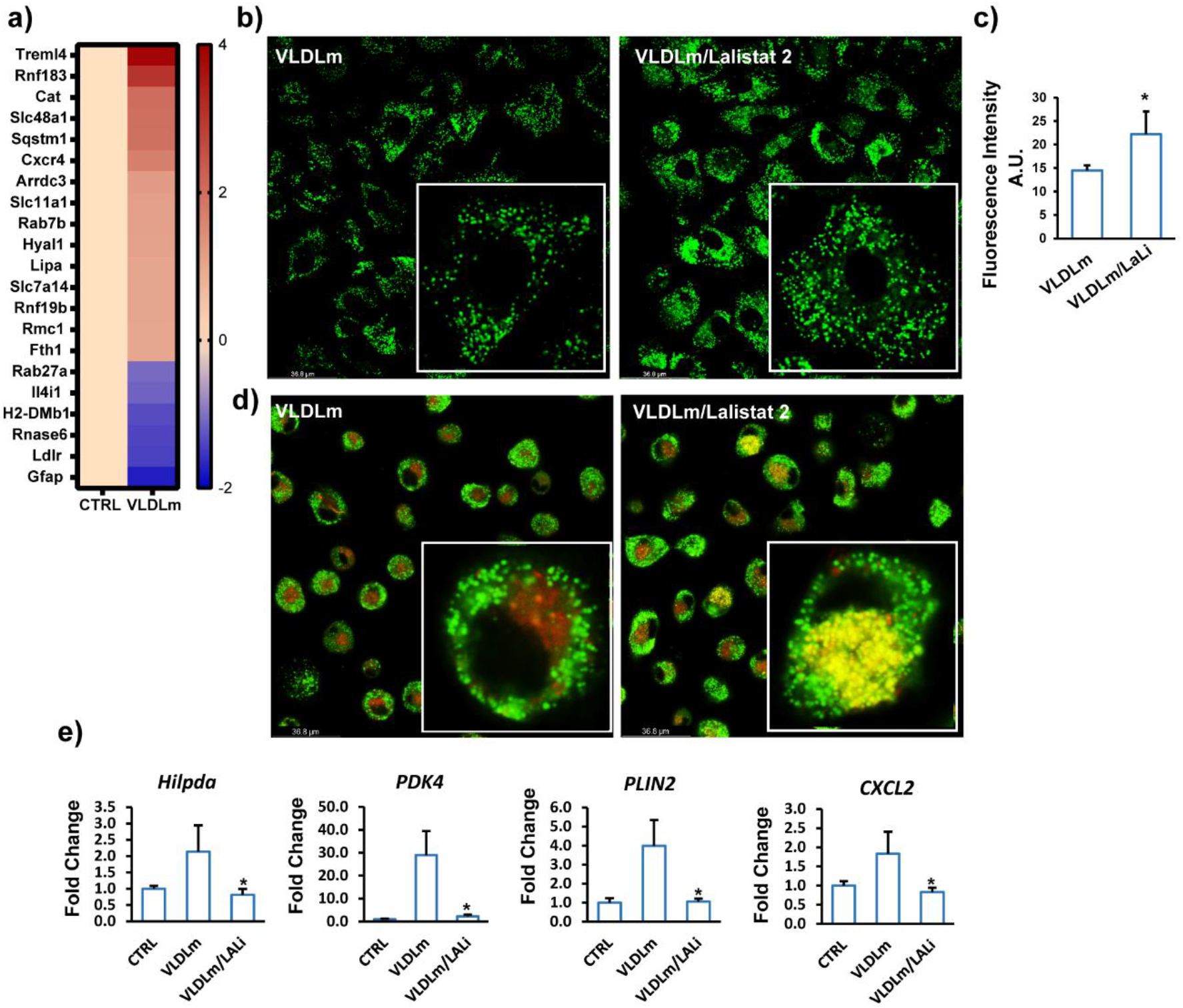
VLDLm-TG are degraded by lysosomal acid lipase. a) Heatmaps showing changes in expression of genes involved in lysosome activity (p<0.01, SLR>1). b) BODIPY 493/503 staining of human macrophages treated with 0.5 mM VLDLm for 6 hours in the presence or absence of 30 μM Lalistat 2. c) Mean fluorescence intensity quantified by flow cytometry. d) Co-staining of lysosome (red) and neutral lipids (green) in human macrophages co-treated with 30 μM Lalistat 2 and 0.5 mM VLDLm for 6 hours. e) mRNA levels of lipid-sensitive genes in human macrophages co-treated with 30 μM Lalistat 2 and 0.5 mM VLDLm for 6 hours.

### NPC1 mediates VLDLm-derived free fatty acid efflux

Inasmuch as NPC1 mediates the lysosomal export of cholesterol, we considered the possibility that NPC1 might also be involved in lysosomal export of VLDL-derived fatty acids. Interestingly, the expression of NPC1 was significantly up-regulated by VLDLm in human primary macrophages (Figure 7a). Co-treatment of VLDLm-treated human primary macrophages with a chemical inhibitor of NPC1 was associated with markedly elevated intracellular lipid accumulation (Figure 7b), as well as with much more pronounced lysosomal staining (Figure 7c). To verify the notion that lipids may be partly retained in the lysosomes upon NPC1 inactivation, we silenced NPC1 in VLDLm-treated human primary macrophages and performed co-staining for lipids and lysosomes. Remarkably, NPC1 silencing (Figure 7d) markedly increased the overlap in lipid and lysosomal staining (Figure 7e), suggesting that NPC1 deficiency causes lipids to be retained in lysosomes. Concurrent with the retention of lipids in the lysosomes, NPC1 silencing significantly decreased the levels of non-esterified fatty acids in the medium of VLDLm-treated macrophages (Figure 7f). Similarly, chemical inhibition of NPC1 significantly blunted the increase in non-esterified fatty acids in the medium of VLDLm-treated macrophages (Figure 7g), concurrent with an increase of lysosome dysfunction-related genes (Figure 7h). These findings suggest that NPC1 promotes the extracellular efflux of fatty acids from lysosomes, which is in agreement with recent data indicating that fatty acids generated by lipophagy can be exported from cells via lysosomal fusion to the plasma membrane [31].

**Figure 7.**
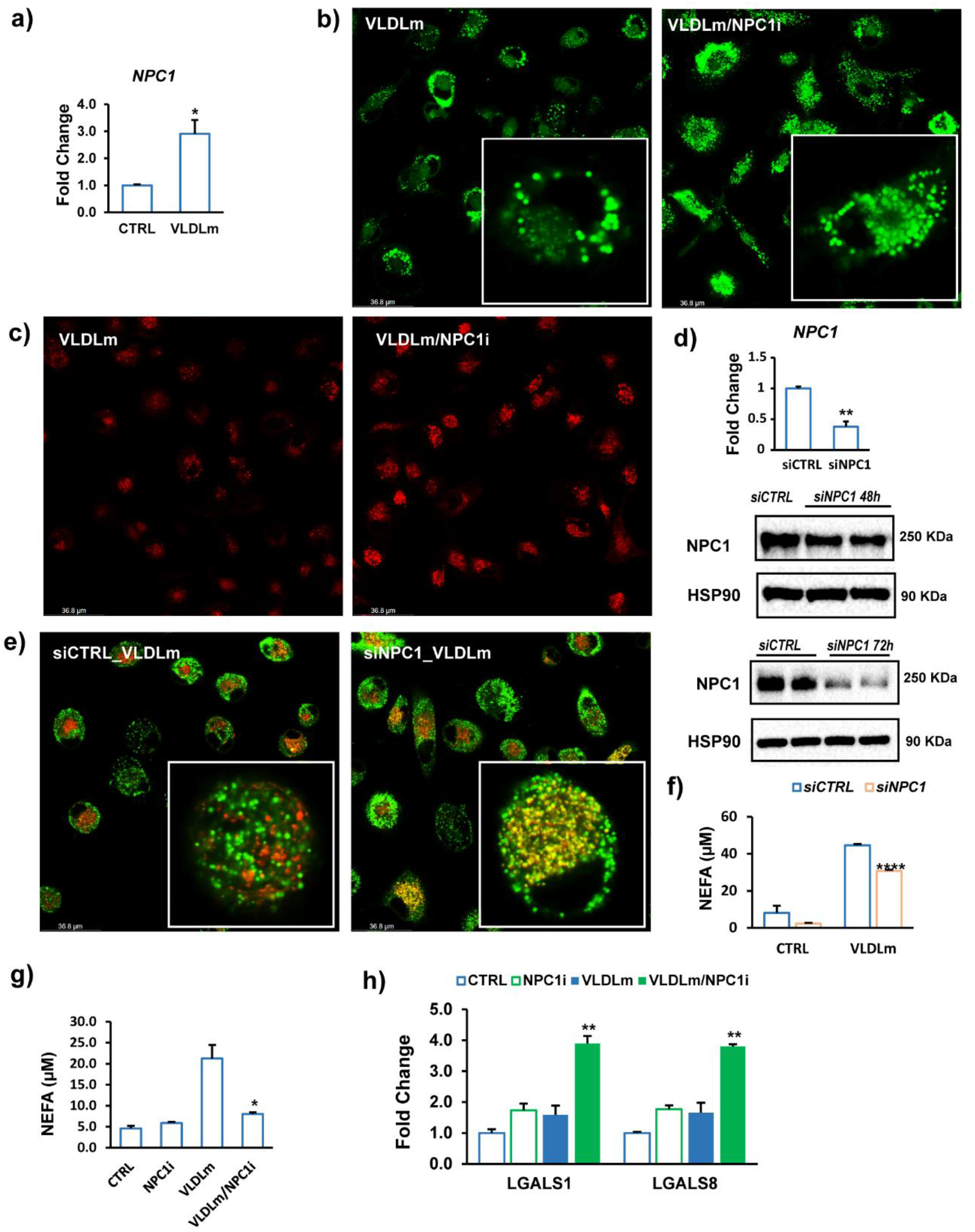
VLDLm-TG are released from macrophages after lysosomal processing. a) mRNA levels of NPC1 in human macrophages. BODIPY 493/503 staining (b) or lysosomal staining (c) of human macrophages treated with 0.5 mM VLDLm for 6 hours in the presence or absence of 5 μM NPC1 inhibitor U18666A. d) mRNA and protein levels of NPC1 in human macrophages treated with siCTRL or siNPC1 e) Co-staining of lysosome (red) and neutral lipids (BODIPY 493/503, green) in human macrophages treated with siCTRL or siNPC1 for 72 hours followed by treatment with 0.5 mM VLDLm for 24 hours and rest overnight. Free fatty acid concentration in culture medium from human macrophages treated with siCTRL or siNPC1 for 48 hours followed by treatment with 0.5 mM VLDLm for 6 hours (f) or treated with 0.5 mM VLDLm for 6 hours in the presence or absence of 5 μM NPC1 inhibitor U18666A (g). h) mRNA levels of Galectin-encoding genes. The bar graphs were plotted as mean±SD. Asterisk indicates significantly different from control according to Student’s t-test. *p<0.05, **p<0.01, ****p<0.001.

### Stard3 is involved in the intracellular transportation of VLDL-derived lipids from the lysosome

In addition to NPC1, the expression of Stard3 was up-regulated by VLDLm in human macrophages (Figure 8a). Inasmuch as Stard3 mediates the transport of cholesterol from lysosome to ER [18], we hypothesized that Stard3 may have a similar role in the intracellular trafficking of VLDLm-derived fatty acids. In line with this hypothesis, silencing of Stard3 (Figure 8b) in VLDLm-treated human macrophages increased lipid accumulation in lysosomes and reduced lipid content in the ER (Figure 8c-d). Unlike silencing of NPC1, silencing of Stard3 did not increase intracellular lipid accumulation nor did it influence efflux of fatty acids into the medium (Figure 8e). These findings suggest that Stard3 is involved in commuting fatty acids from the lysosome to the ER but does not mediate extracellular efflux of fatty acids. Taken together, our data indicate that NPC1 and Stard3 have differential roles in the transport of VLDL-derived fatty acids from the lysosome to other (extra)cellular compartments. Specifically, NPC1 participates in extracellular efflux of lysosomal fatty acids, while Stard3 is involved in transfer of lysosomal fatty acids to the ER for subsequent storage as triglycerides.

**Figure 8.**
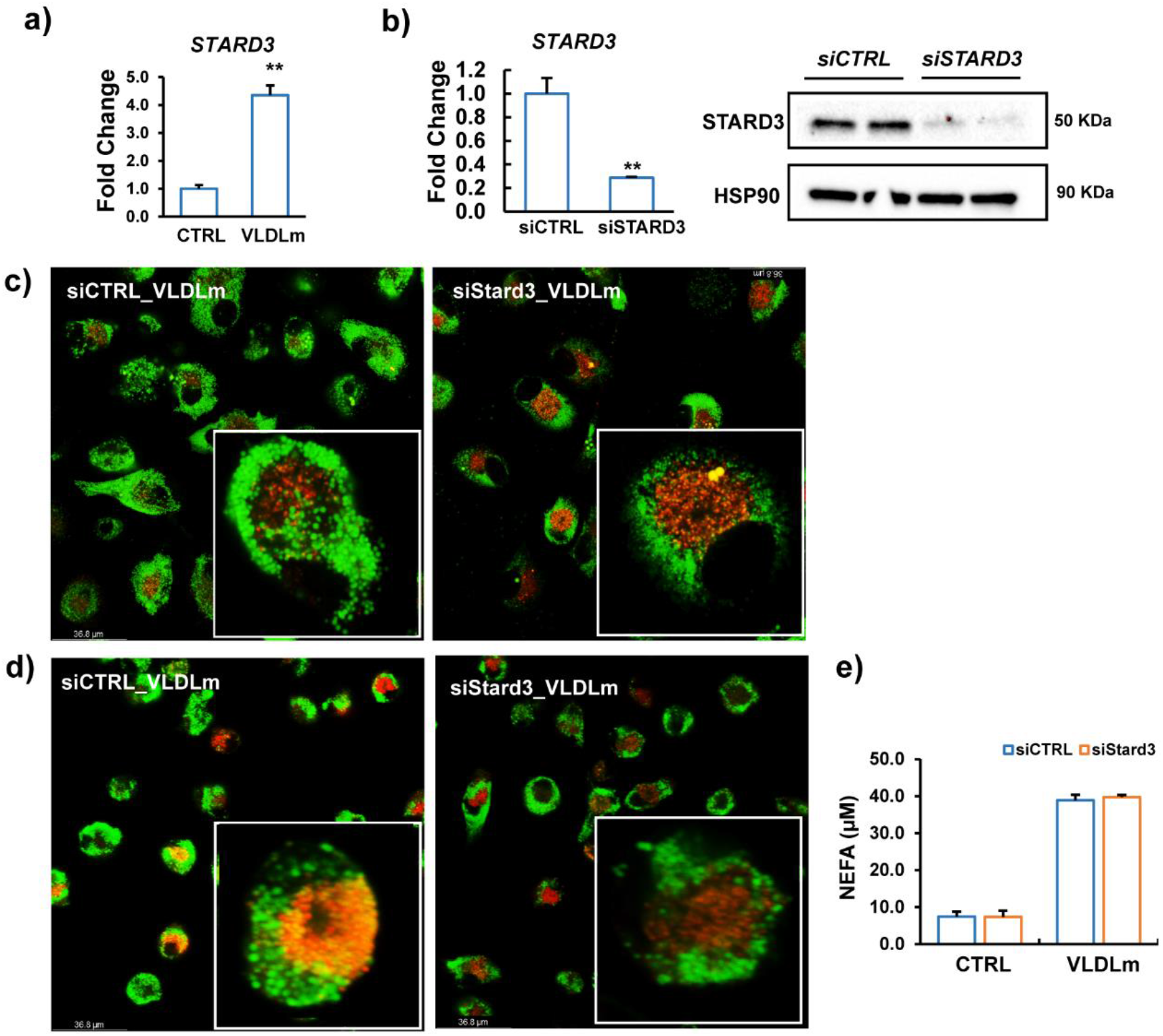
NPC1 and Stard3 have differential roles in the processing of VLDLm-TG. a) mRNA levels of STARD3 in human macrophages. b) mRNA and protein levels of STARD3 in human macrophages treated with siCTRL or siSTARD3 for 24 hours. c) Co-staining of lysosome (red) and neutral lipids (BODIPY 493/503, green) in human macrophages treated with siCTRL or siSTARD3 for 48 hours followed by treatment with 0.5 mM VLDLm for 24 hours and rest overnight. e) Free fatty acid concentration in culture medium from human macrophages treated with siCTRL or siSTARD3 for 24 hours followed by treatment with 0.5 mM VLDLm for 6 hours and rest overnight. The bar graphs were plotted as mean±SD. Asterisk indicates significantly different from control according to Student’s t-test. *p<0.05, **p<0.01, ****p<0.001.

### Macrophages take up chylomicron-like lipoproteins via a similar mechanism as VLDL

In contrast to VLDL, chylomicrons are not able to pass into the intima. Nevertheless, chylomicrons may come into direct contact with macrophages in the mesenteric lymph node. In addition, severely elevated chylomicron levels are associated with the accumulation of chylomicron-derived lipids within skin macrophages, giving rise to eruptive xanthomas. Accordingly, we asked whether the mechanism of uptake of chylomicrons by macrophages resembles the mechanism for uptake of VLDL. To that end, we repeated the studies using lipid-emulsion particles with a mean diameter of 240 nm (Figure S1a), referred to as chylomicron-mimic (CHYLm), in agreement with the reported particle sizes of median-sized chylomicrons[32]. Treatment of RAW 264.7 macrophages and human primary macrophages with these particles for 6 hours led to marked lipid accumulation (Figure S1b) and increased expression of lipid-sensitive genes (Figure S1c). Similar to VLDLm, we found that LPL, but not its catalytic activity, is required for CHYLm uptake by macrophages (Figure S2& Figure S3) and that this process is driven by caveolae-mediated endocytosis (Figure S4 and Figure S5), involving caveolin 1 and 2 (Figure S6). Taken together, these data indicate that the mechanism for macrophage uptake of VLDLm and CHYLm are highly similar.

## Discussion

Due to its ability to enter the intima and be taken up by vascular macrophages, VLDL may contribute the atherosclerosis. Here we studied the mechanism of uptake of VLDL particles by macrophages. We found that macrophage uptake of VLDL requires LPL and is mediated by the lipoprotein-binding C-terminal domain of LPL and not by the catalytic N-terminal domain. Subsequent internalization of VLDL-TG by macrophages was shown to occur via caveolae-mediated endocytosis, followed by triglyceride hydrolysis by LAL in the lysosome. Intriguingly, NPC1 was found to promote the extracellular efflux of fatty acids from lysosomes, while Stard3 is involved in the transfer of lysosomal fatty acids to the ER for subsequent storage as triglycerides. These data suggest that macrophage have the remarkable capacity to excrete part of the internalized triglycerides as fatty acids. Our data elaborate on the model put forward by Lindqvist and colleagues many years ago, who on the basis of studies using radiolabeled and unlabeled human VLDL proposed that incubation of macrophages with VLDL leads to triglyceride accumulation via uptake of intact VLDL, mediated by either a receptor or a nonreceptor-mediated pathway and involving phagocytosis or endocytosis [33].

LPL is mainly known for its ability to catalyze the hydrolysis of TRL-triglycerides and thereby promote the uptake of TRL-derived fatty acids in tissues such as heart, skeletal muscle, white adipose tissue, and brown adipose tissue[32] [34]. In addition to catalyzing TG hydrolysis, LPL can facilitate the binding and uptake of lipoprotein particles by cells independent of lipolysis[35] [36]. Through its ability to interact with lipoproteins on the one hand, and heparan sulfate proteoglycans or specific surface receptors on the other hand, LPL can function as a bridge between lipoproteins and the cell surface [37]. For example, in macrophages—which are characterized by high levels of LPL expression—LPL was found to enhance uptake of oxLDL[38].

Previous studies have shown that LPL promotes the uptake TRL-triglycerides and -cholesterol by macrophages *in vitro* [33] [39]. However, thus far it was not fully clear whether the stimulatory effect of LPL on macrophage uptake of TRL-TG is mainly mediated by the above mentioned lipolysis-independent bridging function—leading to whole particle uptake—or requires actual triglyceride hydrolysis followed by cellular uptake of fatty acids [40]. Using antibodies directed against the N- or C-terminal domain of LPL, we found that uptake of VLDLm-TG by macrophages is dependent on particle-binding by LPL but not on the catalytic function of LPL. Whether binding to HSPG-bound LPL is sufficient to trigger endocytosis of the VLDL-like particles or requires an additional receptor remains to be determined.

Previous evidence suggested that LPL-catalyzed lipolysis is not required for uptake of VLDL-TG by cultured macrophages but does have a modest stimulatory effect [33]. Further work is necessary to better define the role of the catalytic function of LPL in macrophage lipid metabolism. For example, it would be of interest to know the impact of replacement of wildtype LPL by catalytically inactive mutated LPL on lipid uptake and metabolism in macrophages treated with VLDLm emulsion particles [41]. As an alternative, we considered the use of the LPL inhibitor Orlistat. However, a major drawback of orlistat is that it also inhibits LAL [42], which would seriously complicate the interpretation of the data.

Our data suggest that macrophages internalize entire VLDL particles via endocytosis. Uptake of entire TRL particles has been previously observed in vascular endothelial cells of BAT and WAT [43][44][45]. In endothelial cells, the TRL are routed through the endosomal-lysosomal pathway, where they undergo lysosomal acid lipase (LAL)-mediated processing [46]. This route strongly resembles the pathway we observed for VLDL-like particles in human primary macrophages.

It is well established that LDL and oxLDL are taken up by cells via receptor-mediated clathrin-dependent endocytosis [47][48][49][50][51][52]. In contrast, our data indicate that VLDL is taken up by macrophages via caveolae-mediated endocytosis. Specifically, we observed that genistein markedly reduced lipid accumulation in macrophages treated with VLDLm, whereas cholopromazine had no effect. Gene silencing of CAV1 and especially CAV2 also markedly reduced lipid accumulation in VLDL-treated macrophages. The difference in the type of endocytosis for uptake of LDL and VLDL might be explained by the different sizes of the two types of particles or by differences in the composition of the lipid cargo [53].

The internalization and subsequent breakdown of pathogens, apoptotic cells, or particles such as lipoproteins burden the macrophage with potentially toxic macromolecules that must either be metabolized or expelled [54]. For example, internalized cholesterol can either be esterified to fatty acids and stored or it can be exported out of the macrophages via ABCA1 to lipid-poor apolipoprotein A-I to generate precursors for HDL particles. Through these mechanisms, macrophages are able to limit the toxic effects of excessive free cholesterol levels on the cell membrane. Similar to cholesterol, elevated intracellular levels of free fatty acids can also be damaging to cells. To restrict this lipotoxicity, macrophages and other cells are able to convert fatty acids into triglycerides as well as use the fatty acids as fuel. However, in contrast to cholesterol, there is very little evidence in the literature that macrophages export fatty acids. Lindqvist and colleagues showed that the presence of BSA in the culture medium markedly decreased triglyceride content in lipid-laden macrophages, raising the suggestion that macrophages are capable of mobilizing its stored triglycerides [33]. Consistent with these data, we found that macrophages loaded with VLDL-like particles release fatty acids into the medium and additionally we found that this process was impaired by inactivation of the intracellular cholesterol transporter NPC1. At this stage, it is unclear if extracellular fatty acid efflux is dependent on the fusion of the lysosomes to the plasma membrane or requires fatty acid transport through the cytoplasm. Since NPC1 was specifically dismissed as fatty acid transporter [55], the decreased fatty acid release upon NPC1 inactivation is probably secondary to impaired lysosomal lipolysis due to lysosomal accumulation of cholesterol, rather than reflecting a direct role of NPC1 as fatty acid transporter.

Consistent with our data, Mashek and colleagues recently found that fatty acids liberated by lipophagy in lysosomes can be transported out of the cell, and suggested that the effluxed fatty acids may be available for uptake by the same cell[31]. Based on our data, it is impossible to say whether the extracellular efflux of lysosome-derived fatty acids is needed for the storage of VLDL-TG in macrophages via re-uptake of the fatty acids. One could wonder about the rationale for exporting fatty acids from the cell if most of the fatty acids are re-taken up. Rather, the efflux of fatty acid may be a mechanism to rid the macrophage of excess fatty acids acquired via endocytosis and phagocytosis. It can be speculated that the fatty acids released by macrophages may be used as fuel by neighboring cells.

StAR-related lipid transfer domain-3 (Stard3) is a sterol-binding protein that promotes sterol transport by creating endoplasmic reticulum (ER)–endosome contact sites [56]. We found that Stard3 is involved in the transfer of VLDL-derived fatty acids from the lyso/endosome to the ER. Whether Stard3 directly binds fatty acids or promotes transport by creating membrane contact sites requires further study. Our data suggest that in contrast to NPC1, Stard3 does not mediate the extracellular efflux of fatty acids. NPC1 and Stard3 thus have distinct roles in the transport of VLDL-derived fatty acids from the lysosome to other (extra)cellular compartments.

Our key findings on VLDLm particle uptake and metabolism in macrophages were reproduced when using larger CHYLm particles, suggesting that TRL are processed through a single mechanism. However, it should be noted that the range in particle sizes of our artificial CHYLm particles is smaller than that of human chylomicrons. Accordingly, it cannot be excluded that human chylomicrons are also taken up via other mechanisms, such as micropinocytosis. Previous *in vitro* studies already found large similarities in the mechanism of macrophages uptake and degradation between VLDL and chylomicrons, showing among other things that macrophages are able to take up intact VLDL and chylomicron particles [33][39]. The *in vivo* relevance of uptake of chylomicrons by macrophages is likely limited. In contrast to VLDL and its remnants, chylomicrons are not able to penetrate the arterial wall to be taken up by macrophages. One specific location where chylomicrons do get in direct contact with macrophages is in the mesenteric lymph nodes, where excessive uptake of chylomicrons can lead to formation of giant macrophage foam cells, as observed in ANGPTL4 deficient mice on a high fat diet [57].

Our study also has limitations. First, most of our experiments were conducted with artificial triglyceride-rich emulsion particles. These VLDL-mimicking particles were generated with a microfluidizer using casein as emulsifier. It should be noted, though, that our key findings were verified with human VLDL. In addition, our data on the role of LPL in TG uptake are fully consistent with previous studies that used human VLDL. The similarity in the mechanism of uptake of human VLDL and artificial VLDLm particles containing casein as emulsifier suggest that macrophages recognize TRL particles on the basis of size and/or lipid content rather than a specific protein component. A second limitation of our study is that the triglycerides in the VLDLm particles were not radioactively or fluorescently labeled. Consequently, we cannot fully exclude that part of the stored lipids may be endogenously synthesized by the macrophage in response to treatment with VLDLm. However, this contribution likely is small. In the future, fluorescently-labeled triglycerides could be incorporated into the VLDLm particles to enable direct intracellular visualization of exogenous lipids.

In conclusion, our data suggest that the uptake of TRL-derived triglycerides by macrophages is mediated by the lipid-binding function of LPL. After binding to LPL, TRL-derived triglycerides are taken up by caveolin-mediated endocytosis, followed by LAL-catalyzed hydrolysis in the lysosomes. Subsequent processing of TRL-derived fatty acids towards storage requires the proteins NPC1, which was found to promote the extracellular efflux of fatty acids from lysosomes, and Stard3, which is involved in the transfer of lysosomal fatty acids to the ER for subsequent storage as triglycerides. Our data provide key new insights into how macrophages take up and process triglyceride-rich lipoproteins.

## Methods

### Preparation of VLDL- and chylomicron-sized lipid emulsions

The VLDL- and chylomicron-sized lipoprotein emulsions (referred as VLDLm and CHYLm respectively) were prepared with commercial sunflower oil (Gwoon, the Netherlands) with a microfluidizer at pressures of 400 bar and 1200 bar, respectively, for five cycles. Sodium caseinate (purity 97%, Excellion, FrieslandCampina, the Netherlands) was used as emulsifier. The size of the droplets was measured using a Mastersizer 3000 (Malvern Panalytical, United Kingdom) and presented in d (3,2).

### Cell culture

#### RAW 264.7 macrophages

RAW 264.7 macrophages were cultured in DMEM supplemented with 10% FCS and 1% p/s. Cells were seeded at the density of 52,000 cells/ cm^2^ and cultured overnight before treatment with emulsions for 6 hours.

### Human buffy-coat primary macrophages

#### PBMC isolation

Human buffy-coat blood was obtained from Sanquin, the Netherlands. Briefly, 25 ml of buffy-coat blood diluted 1:1 with PBS was added to 50 ml Leucosept tubes containing 15 ml of Ficoll-Paque followed by centrifugation for 15 min at 800 RCF at room temperature. Afterwards, the PBMC layer was collected and washed with cold PBS for three times. 70 μm cell strainers were used to remove clumps/ fat residues.

#### Monocytes isolation

Moncytes were isolated by MojoSort CD14 positive selection kits (Biolegend, California, United States) and LS columns (Miltenyi Biotec, Bergisch Gladbach, Germany). Briefly, 10 μl CD14 Nanobeads (10x pre-diluted in MACS buffer (PBS+ 0.5%BSA+ 2mM EDTA)) and 90 μl MACS buffer were added per 1×10^7^ PBMCs and incubated at 4°C for 15 min (gently mixed every 5 min). Afterwards, the beads conjunct cells were separated by LS columns using a Miltenyi QuadroMACS Separator.

Isolated cells were cultured in RPMI medium supplemented with 10% FCS, 1% GlutaMax and 1% p/s at a density of 1×10^6^ cells/ml. 5 ng/ml of Granulocyte-macrophage colony-stimulating factor (GM-CSF; Miltenyi) was used to differentiate monocytes into macrophages. After full differentiation, cells were seeded at the density of 100,000 cells per cm^2^ for siRNA assay and 150,000 cells per cm^2^ for the other assays.

#### Inhibitors and heparin assay

Cells were pre-incubated with chemical inhibitors (30 μM of Lalistat 2, 0.2 μM of GSK264220A, 200 μM of Genistein, 10 μg/ml of chlorpromazine, 5 μM of U18666A) for 1 hour and then continuously treated with lipid emulsions for 6 hours.

For the heparin assay, cells were treated with 50 UI/ml of heparin for 2 hours, followed by two times washing with PBS. Emulsions were subsequently added to the cells together with the same concentration of heparin.

Human LPL F:1 (Santa Cruz Biotechnology, Inc., California, USA) and 5D2 antibodies were added to cell culture medium 2 hours prior to emulsion loading at the concentration of 2 μg/ml (1:100 dilution).

Genistein, Lalistat 2, chlorpromazine and heparin were obtained from Sigma-Aldrich, Missouri, United States. GSK264220A was from Tocris (bio-techne), Abingdon, United Kingdom. LPL 5D2 antibody was contributed by Department of Medicine, David Geffen School of Medicine, University of California, Los Angeles, USA.

#### Free fatty acids assay

For functional study of NPC1 and Stard3, GM-CSF derived macrophages were loaded with artificial VLDL particles. Afterwards, the medium was refreshed and cells were rested overnight. Then medium was collected for assessment of free fatty acids using the free fatty acids kit (Instruchemi, The Netherlands) following manufacturer’s instructions.

#### Quantitive RT-PCR

Total RNA was isolated using TRizol ™ Reagent (Thermo Fisher Scientific, Massachusetts, United States). cDNA was synthesized using iScript ™ cDNA Sythesis Kit (Bio-Rad Laboratories, Inc., California, United States) following the manufacturer’s protocol. Real-Time polymerase chain reaction (RT-PCR) was performed on the CFX 384 Touch™ Real-Time detection system (Bio-Rad Laboratories, Inc., California, United States), using the SensiMix ™ (BioLine, London, UK) protocol for SYBR green reactions. Mouse/human 36B4 expression was used for normalization.

#### Immunoblotting

The cell lysates were prepared using RIPA Lysis and Extraction Buffer (Thermo Fisher Scientific, Massachusetts, United States) or with self-prepared NP40 lysis buffer (50 mM Tris-HCl pH 8.0, 0.5% NP40, 150 mM NaCl, 5 mM MgCl_2_) for cell membrane binding proteins (LPL, Caveolin 1) and quantified with Pierce™ BCA Protein Assay Kit (Buffer (Thermo Fisher Scientific, Massachusetts, United States). The cell lysates were separated by electrophoresis on pre-cast 4-15% polyacrylamide gels and transferred onto nitrocellulose membranes using a Trans-Blot® Semi-Dry transfer cell (Bio-Rad Laboratories, Inc., California, United States), blocked in 5% skim milk in TBS-T (TBS buffer supplied with 1 ‰ TWEEN 20) and incubated with LPL antibody (F:1), caveolin-1 Antibody (4H312), Stard3 antibody (H-1) (sc-166215) (Santa Cruz Biotechnology, Inc., Texas, United States) and NPC1 antibody (ab134113) (Abcam, Abcam, Cambridge, United Kingdom) overnight at 4°C. Secondary antibody incubation was performed at room temperature for 1 hour. Hsp90 was used for normalization (antibody was purchased from Cell Signaling Technology, Inc., Massachusetts, United States). Images were gained using the ChemiDoc MP system (Bio-Rad Laboratories, Inc., California, United States).

#### Transcriptomics

Cells were treated with 0.5 mM of lipoprotein mimic emulsions for 6 hours and collected for total RNA isolation. The experiments were performed in both biological and technical triplicates. Samples of each condition from one biological study were pooled. Transcriptome analysis by RNA-sequencing was performed by BGI Hong Kong Company Limited (Hong Kong) following a standard protocol. In brief, RNA samples were prepared using RNeasy Mini Kit (Qiagen, Hilden, Germany) following manufacturer’s instructions. mRNA molecules were purified from total RNA using oligo(dT)-attached magnetic beads and fragmented into small pieces using fragmentation reagent after reaction for a certain period at proper temperature. First-strand cDNA was generated using random hexamer-primed reverse transcription, followed by a second-strand cDNA synthesis. The synthesized cDNA was subjected to end-repair and then was 3’ adenylated. Adapters were ligated to the ends of these 3’ adenylated cDNA fragments. PCR products were purified with Ampure XP Beads (AGENCOURT) and dissolved in EB solution. Library was validated on the Agilent Technologies 2100 bioanalyzer. The double stranded PCR products were heat denatured and circularized by the splint oligo sequence. The single strand circle DNA (ssCir DNA) were formatted as the final library. The library was amplified with phi29 to make DNA nanoball (DNB) which had more than 300 copies of one molecule. The DNBs were load into the patterned nanoarray and pair end 100 or 150 bases reads were generated in the way of sequenced by synthesis.

#### Confocal imaging

Cells were seeded and treated in μ-Slide 8 Well Glass plates (ibidi GmbH, Planegg, Germany) and visualized with Leica SP8-SMD confocal microscope (Leica Microsystems, Wetzlar, Germany) equipped with a 63× 1.20 NA water-immersion objective lens. Images were acquired using 1,024 × 1,024 pixels with the pinhole set at 1 Airy Unit (AU). Excitation of the fluorescent probes was performed using white light laser (WLL, 50% laser output). Florescent emission was detected using an internal Hybrid (HyD) detector.

Lipid droplet accumulation was measured on 3.7% formaldehyde fixed cells after 20 min incubation with 2μg/ml BODIPY™ 493/503 (Thermo Fisher Scientific, Massachusetts, United States) and mounted with Vectashield-H anti-fade medium (Vector laboratories, California USA). The WLL laser line (488 nm) was set at a laser power of 2.5% and emission was detected selecting a spectral window of 505-550 nm.

Endoplasmic reticulum (ER) was stained in live cells using ER Staining Kit - Red Fluorescence - Cytopainter (ab139482) (Abcam, Cambridge, United Kingdom) following the manufacturer’s protocol. Briefly, cells were stained with 1.5 μl/ml Detection Reagent and cultured with colorless DMEM for 45 minutes at 37°C before detection by microscopy. The WLL laser line (596 nm) was set at a laser power of 5% and emission was detected selecting a spectral window of 670-720 nm.

Lysosome were stained in live cells using LysoTracker™ Deep Red (Thermo Fisher Scientific, Massachusetts, United States) following the manufacturer’s protocol. Briefly, cells were stained with 75 nM fluorescence probe and cultured with colorless DMEM for 60 minutes at 37°C before detection by microscopy. The WLL laser line (647 nm) was set at a laser power of 5% and emission was detected selecting a spectral window of 670-720 nm.

Early endosomes were stained by CellLight™ Early Endosomes-RFP, BacMam 2.0 Kit (Thermo Fisher Scientific, Massachusetts, United States). In brief, cells were incubated overnight with 40 μl/ml CellLight® reagent, fixed with 3.7% formaldehyde and mounted with Vectashield-H anti-fade medium. The WLL laser line (555 nm) was set at a laser power of 5% and emission was detected selecting a spectral window of 565-700 nm. Human primary macrophages for co-staining assays were cultured overnight in fresh medium after the lipid treatment to allow for sufficient intracellular lipid transportation.

#### Flow cytometry analysis

Human primary macrophages (GM-CSF) were seeded in 24-well plates with a density as previously described and treated with 0.5 mM lipoprotein mimic emulsions for 6 hours. 1 μg/ml BODIPY 493/503™ was used for cellular lipid droplet staining. After 20 minutes of incubation with BODIPY at 37°C, cells were twice washed with PBS and then trypsinized. Samples were measured on a CytoFLEX cytometer (Beckman Coulter, Inc, Indianapolis, USA) and data were analyzed by FlowJo (BD, Oregon, USA).

#### siRNA gene knock down assay

Silencing of *LPL, CAV1, CAV2, NPC1* and *STARD3* in GM-CSF buffy coat human macrophages was carried out using ON-TARGETplus siRNA SMARTpool kits (Horizon Discovery Research company, Waterbeach, United Kingdom) following the instructions of manufacturer. Briefly, GM-CSF macrophages were seeded in the desired plates at the density mentioned before and cultured overnight. 50 nM of siRNA was applied together with Lipofectamine™ RNAiMAX Transfection Reagent (Thermo Fisher Scientific, Massachusetts, United States) for 48 hours. Real time-qPCR and immunoblotting were used to assess the transfection efficiency.

#### Statistical Analysis

Data are presented as mean ±SD. Data analysis were performed using unpaired Student’s *t* test. GO Cellular components analysis was completed by Enrichr[58] [59] [60]. All other plots on transcriptome data were generated by Graphpad Prism 8.

## Acknowledgements

Funding from FrieslandCampina (Pacman) is gratefully acknowledged. The authors would like to thank Chang Sun for the help of screening chemical inhibitors of endocytosis, Montserrat A. de la Rosa Rodriguez and Qi Zheng for the help with confocal microscope technology, and Benthe van der Lugt for helping with human monocytes isolation.

## Supplementary materials

**Fig. S1.**
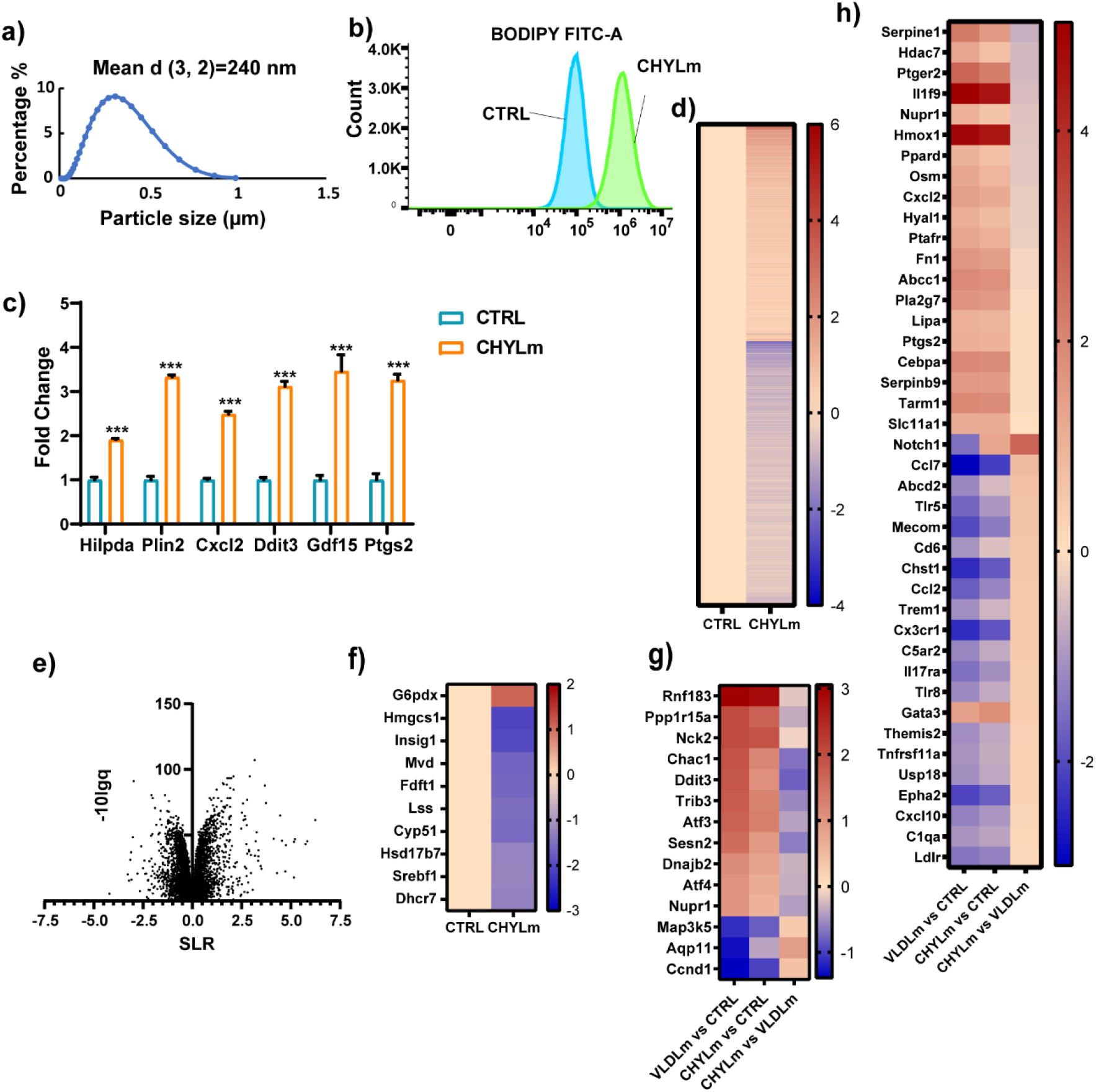
CHYLm promotes lipid accumulation in cultured macrophages. a) The particle size distribution of CHYLm as determined by mastersizer 3000. b) Mean fluoresce intensity (FITC-A) measured by flow cytometry of mouse RAW 264.7 macrophages treated with 1 mM CHYLm for 6 hours. c) mRNA expression of lipotoxic marker genes in RAW 264.7 macrophages treated with 1mM CHYLm for 6 hours. d) Heatmap and e) Volcano plot of RNAseq data of RAW 264.7 macrophages treated with 1 mM CHYLm for 6 hours. f) Heatmap plotted with differentially expressed genes (p<0.01, SLR>1) involved in the cholesterol synthesis pathway. g) Heatmap plotted with differentially expressed genes (p<0.01, SLR>1) Involved in ER stress inflammatory response pathways. Bar graphs were plotted as mean ± SD. Significance was analysed using Two-way ANOVA; *p < 0.05, **p<0.01, ***p<0.001, ***p<0.0001.

**Fig. S2.**
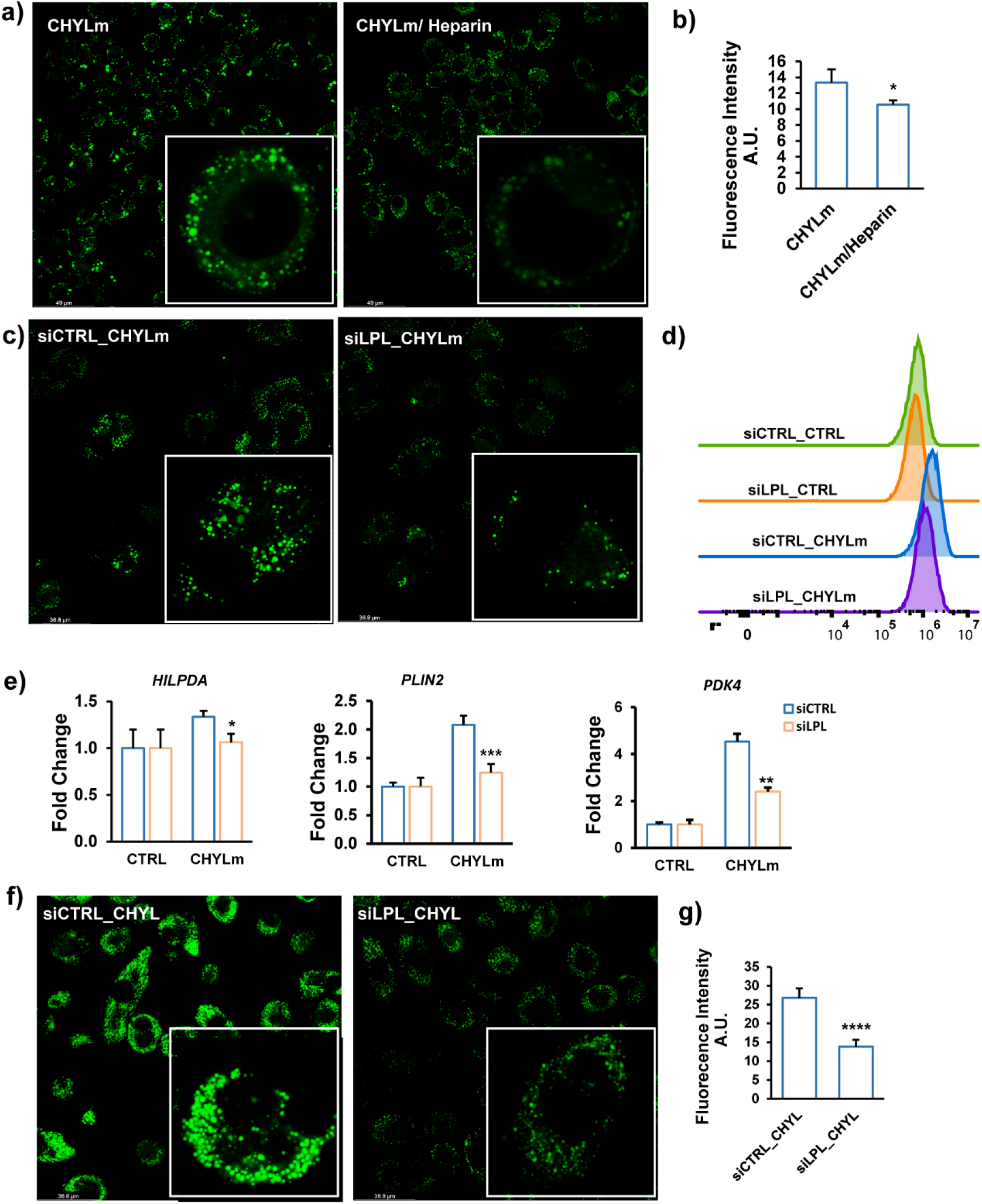
LPL mediates CHYLm uptake in cultured macrophages. a) BODIPY 493/503 staining of intracellular neutral lipids in RAW 264.7 macrophages treated with 1mM CHYLm for 6 hours in the presence or absence of 50 UI/ ml human heparin. b) Quantification of the fluorescence images by ImageJ. c) BODIPY 493/503 staining of intracellular neutral lipids in human macrophages treated with siCTRL or siLPL for 48 hours followed by treatment with 0.5 mM CHYLm for 6 hours. d) Mean fluorescence intensity quantified by flow cytometry. e) mRNA expression of selected lipid-sensitive genes. f) BODIPY 493/503 staining of intracellular neutral lipids in human macrophages treated with siCTRL or siLPL for 48 hours followed by treatment with 0.5 mM human plasma isolated CHYL for 6 hours. g) Mean fluorescence intensity quantified by flow cytometry. The bar graphs were plotted as mean±SD. Asterisk indicates significantly different from control according to Student’s t-test. *p<0.05, **p<0.01, ***p<0.001.

**Fig. S3.**
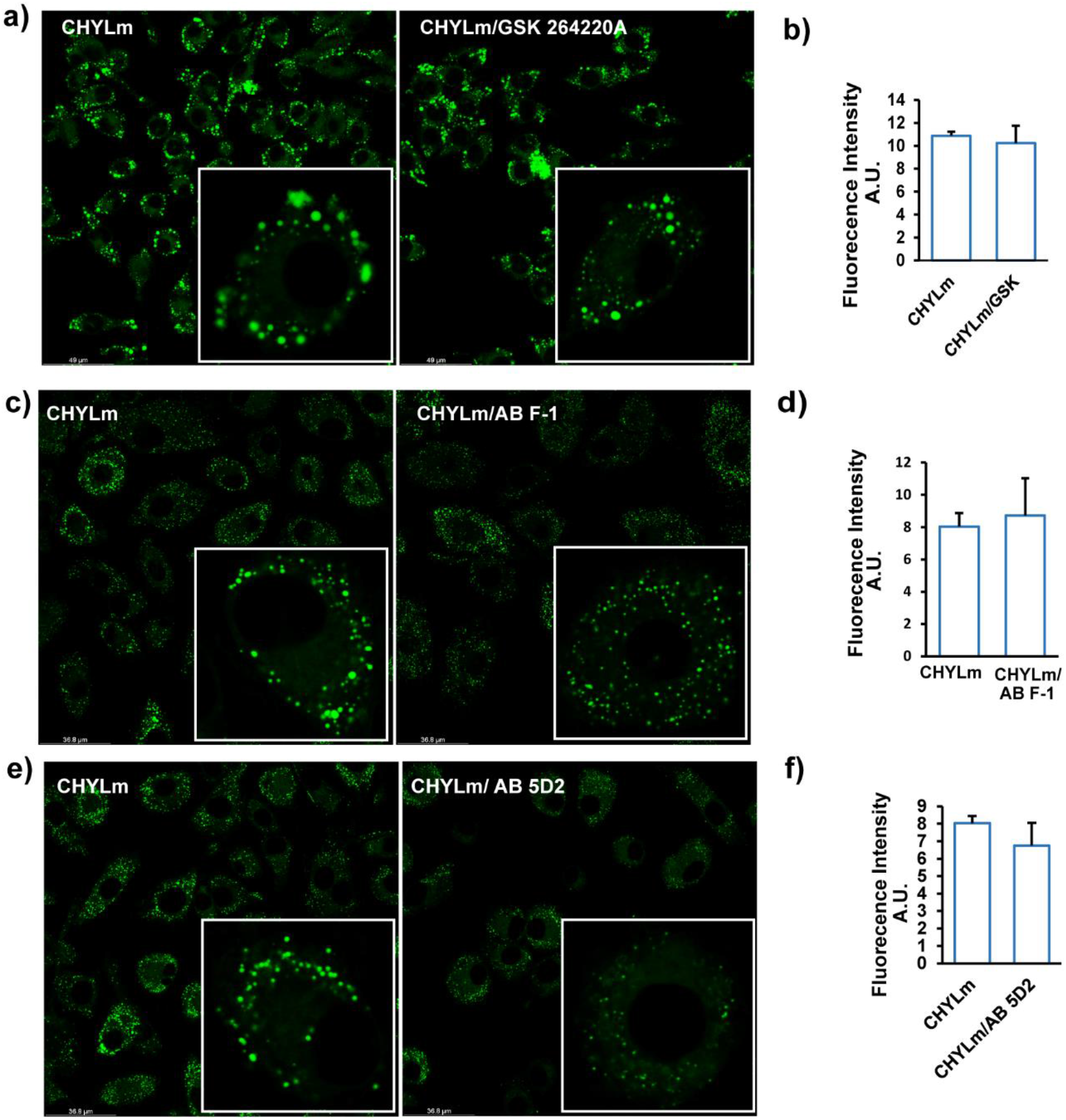
The c-terminal portion of LPL mediates CHYLm uptake in cultured macrophages. a) BODIPY 493/503 staining of RAW 264.7 macrophages treated with 1mM CHYLm for 6 hours in the presence or absence of 0.2 μM of the catalytic LPL inhibitor GSK264220. b) Mean fluorescence intensity quantified by flow cytometry. c) BODIPY 493/503 staining of RAW 264.7 macrophages treated with 1mM CHYLm for 6 hours in the presence or absence of antibody F1 targeting the N-terminal portion of LPL (2 μg/ml). d) Mean fluorescence intensity quantified by Image J. e) BODIPY 493/503 staining of RAW 264.7 macrophages treated with 1mM CHYLm for 6 hours in the presence or absence of antibody 5D2 targeting the C-terminal of LPL (2 μg/ml), f) Mean fluorescence intensity quantified by ImageJ. The bar graphs were plotted as mean±SD. Asterisk indicates significantly different from control according to Student’s t-test. ****p<0.0001.

**Fig. S4.**
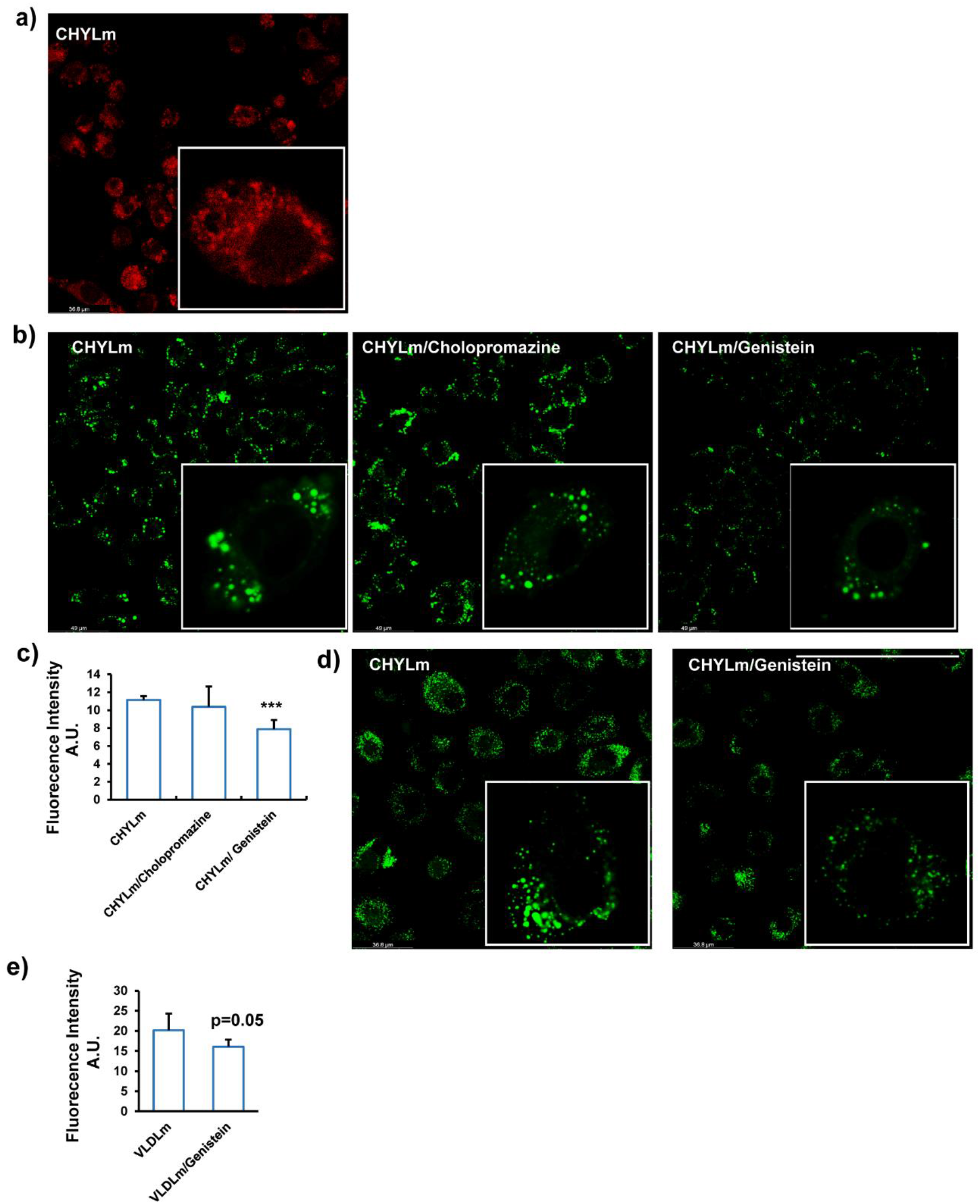
CHYLm are taken up by macrophages via caveola-mediated endocytosis. a) Early endosome staining of RAW 264.7 macrophages treated with 1 mM treatment of CHYLm for 6 hours. b) BODIPY 493/503 staining of RAW 264.7 macrophages treated with 1mM CHYLm for 6 hours in the presence or absence of 10μg/ml chlorpromazine or 200 μM Genistein. c) Mean fluorescence intensity quantified by Image J. d) BODIPY 493/503 staining of human primary macrophages treated with 0.5 mM CHYLm for 6 hours in the presence or absence of 200 μM Genistein. e) Mean fluorescence intensity quantified by ImageJ. The bar graphs were plotted as mean±SD. ***p<0.001.

**Fig. S5.**
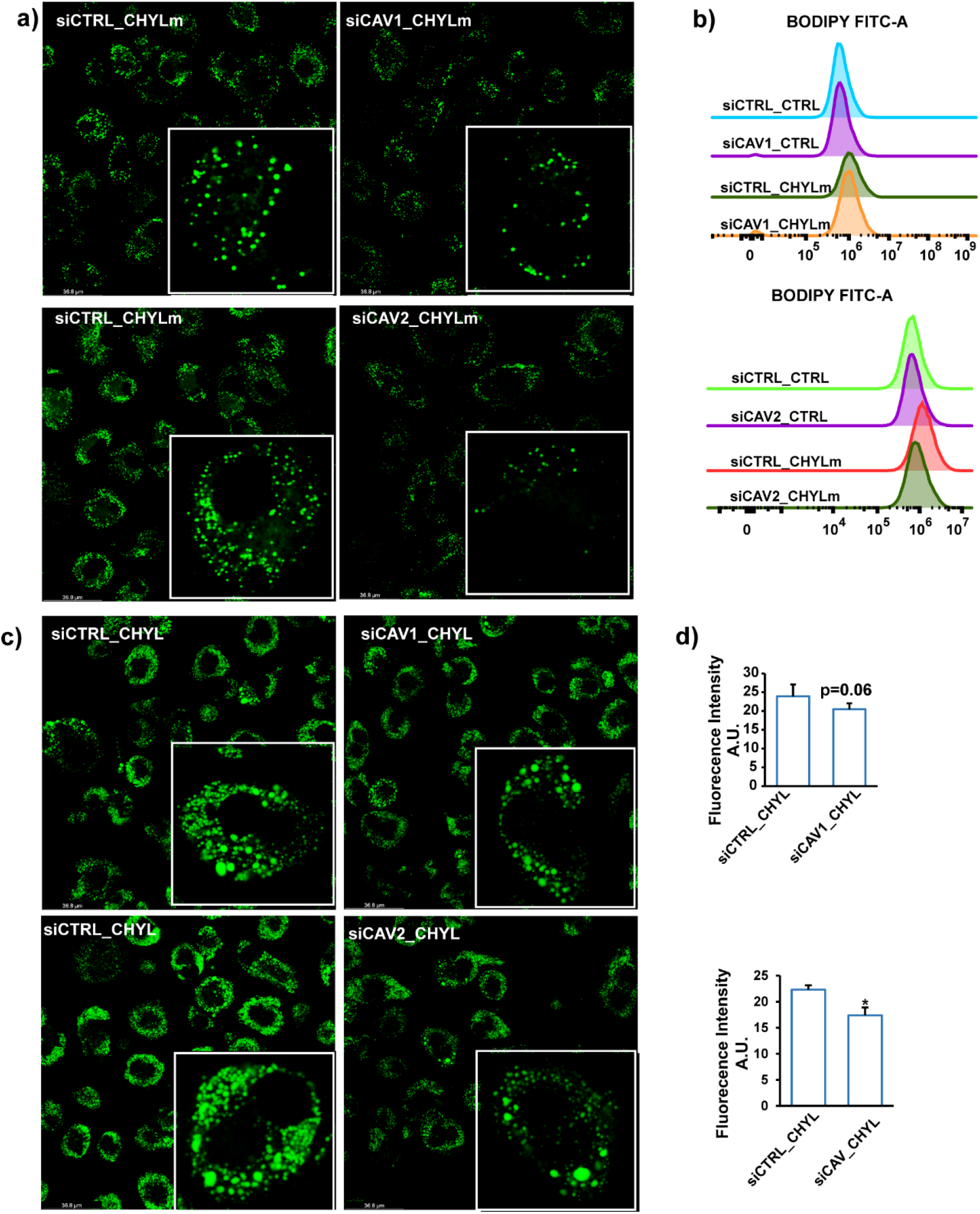
Silencing of Caveolin 1 and 2 impairs uptake of CHYLm by macrophages. a) BODIPY 493/503 staining of human macrophages treated with siCTRL, siCAV1, or siCAV2 for 48 hours followed by treatment with 0.5 mM CHYLm for 6 hours. b) Mean fluorescence intensity quantified by flow cytometry. c) BODIPY 493/503 staining of human macrophages treated with siCTRL, siCAV1, or siCAV2 for 48 hours followed by treatment with 0.5 mM human plasma isolated CHYL for 6 hours. d) Mean fluorescence intensity quantified by ImageJ. The bar graphs were plotted as mean±SD. Asterisk indicates significantly different from control according to Student’s t-test. *p<0.05.

**Fig. S6.**
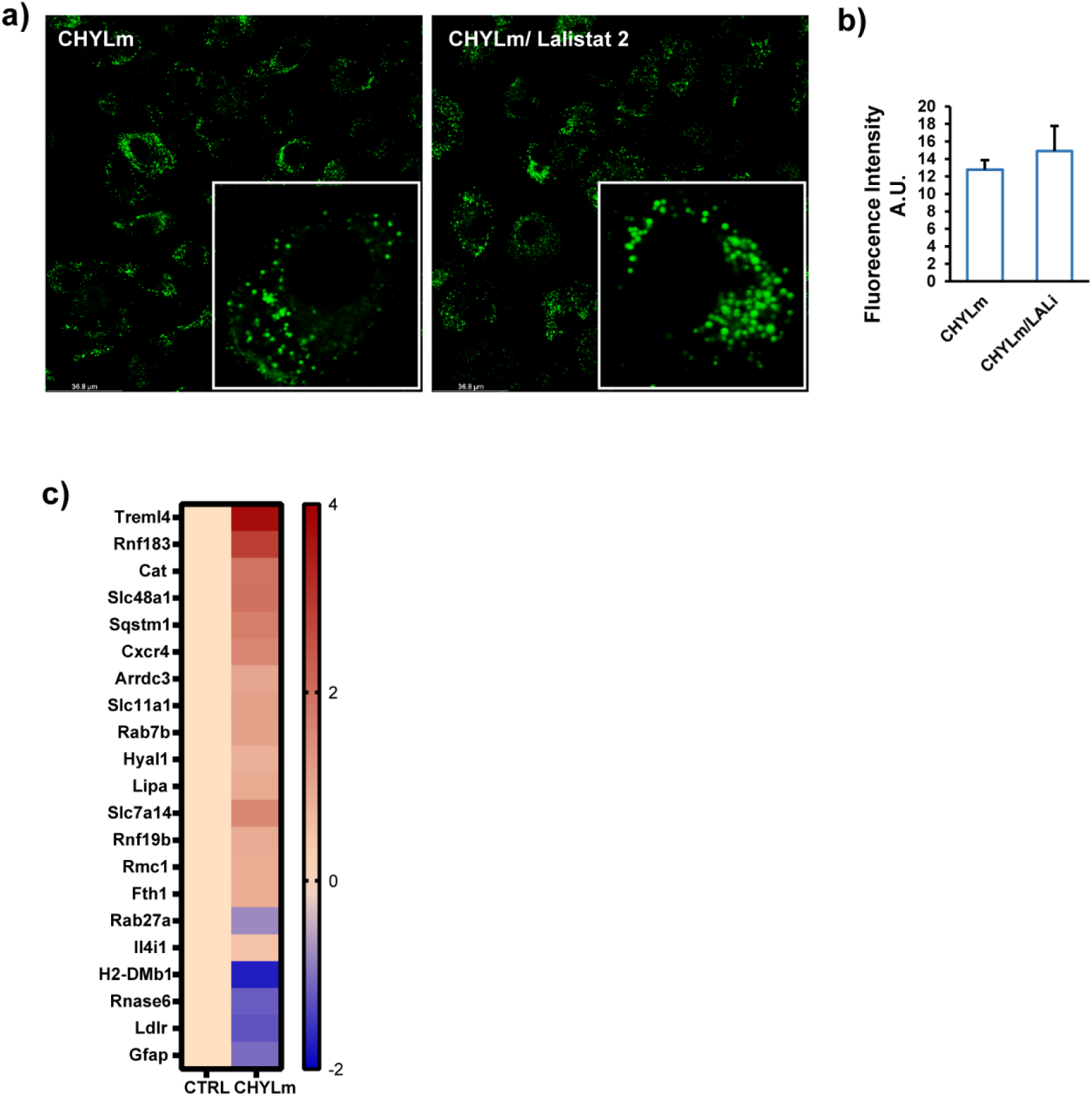
CHYLm-TG are degraded by lysosomal acid lipase. a) BODIPY 493/503 staining of human macrophages treated with 0.5 mM CHYLm for 6 hours in the presence or absence of 30 μM Lalistat 2. b) Mean fluorescence intensity quantified by flow cytometry. d) Co-staining of lysosome (red) and neutral lipids (green) in human macrophages co-treated with 30 μM Lalistat 2 and 0.5 mM CHYLm for 6 hours. c) Heatmaps showing changes in expression of genes involved in lysosome activity (p<0.01, SLR>1).

## References

1. Martins IJ, Mortimer B, Miller J, Redgrave TG. Effects of particle size and number on the plasma clearance of chylomicrons and remnants. J Lipid Res. 1996;37. doi:10.1016/S0022-2275(20)37472-1

2. German JB, Smilowitz JT, Zivkovic AM. Lipoproteins: When size really matters. Curr Opin Colloid Interface Sci. 2006;11: 171. doi:10.1016/J.COCIS.2005.11.006

3. de Gaetano M, Crean D, Barry M, Belton O. M1- and M2-type macrophage responses are predictive of adverse outcomes in human atherosclerosis. Front Immunol. 2016;7: 275. doi:10.3389/fimmu.2016.00275

4. Poznyak A V., Nikiforov NG, Markin AM, Kashirskikh DA, Myasoedova VA, Gerasimova E V., et al. Overview of OxLDL and Its Impact on Cardiovascular Health: Focus on Atherosclerosis. Front Pharmacol. 2020;11. doi:10.3389/FPHAR.2020.613780

5. Saraswathi V, Hasty AH. The role of lipolysis in mediating the proinflammatory effects of very low density lipoproteins in mouse peritoneal macrophages. J Lipid Res. 2006;47: 1406–1415. doi:10.1194/jlr.M600159-JLR200

6. Nordestgaard BG, Wootton R, Lewis B. Selective Retention of VLDL, IDL, and LDL in the Arterial Intima of Genetically Hyperlipidemic Rabbits In Vivo. Arterioscler Thromb Vasc Biol. 1995;15: 534–542. doi:10.1161/01.ATV.15.4.534

7. Dhaliwal BS, Steinbrecher UP. Scavenger receptors and oxidized low density lipoproteins. Clinica Chimica Acta. Clin Chim Acta; 1999. pp. 191–205. doi:10.1016/S0009-8981(99)00101-1

8. Chang HR, Josefs T, Scerbo D, Gumaste N, Hu Y, Huggins LA, et al. Role of LpL (Lipoprotein Lipase) in macrophage polarization in vitro and in vivo. Arterioscler Thromb Vasc Biol. 2019;39: 1967–1985. doi:10.1161/ATVBAHA.119.312389

9. Beisiegel U. New aspects on the role of plasma lipases in lipoprotein catabolism and atherosclerosis. Atherosclerosis. Elsevier Ireland Ltd; 1996. pp. 1–8. doi:10.1016/0021-9150(95)05792-7

10. Ishibashi S, Yamada N, Shimano H, Mori N, Mokuno H, Gotohda T, et al. Apolipoprotein E and lipoprotein lipase secreted from human monocyte-derived macrophages modulate very low density lipoprotein uptake. J Biol Chem. 1990;265: 3040–3047. Available: https://pubmed.ncbi.nlm.nih.gov/2303437/

11. Wendland B. Everything you ever wanted to know about endocytosis. Nat Cell Biol. 2001;3: E254–E254. doi:10.1038/ncb1101-e254

12. Lakadamyali M, Rust MJ, Zhuang X. Ligands for clathrin-mediated endocytosis are differentially sorted into distinct populations of early endosomes. Cell. 2006;124: 997–1009. doi:10.1016/j.cell.2005.12.038

13. Wilhelm LP, Wendling C, Védie B, Kobayashi T, Chenard M, Tomasetto C, et al. STARD 3 mediates endoplasmic reticulum-to-endosome cholesterol transport at membrane contact sites. EMBO J. 2017;36: 1412–1433. doi:10.15252/embj.201695917

14. Zanoni P, Velagapudi S, Yalcinkaya M, Rohrer L, von Eckardstein A. Endocytosis of lipoproteins. Atherosclerosis. Elsevier Ireland Ltd; 2018. pp. 273–295. doi:10.1016/j.atherosclerosis.2018.06.881

15. Mesmin B, Antonny B, Drin G. Insights into the mechanisms of sterol transport between organelles. Cellular and Molecular Life Sciences. Cell Mol Life Sci; 2013. pp. 3405–3421. doi:10.1007/s00018-012-1247-3

16. Infante RE, Wang ML, Radhakrishnan A, Hyock JK, Brown MS, Goldstein JL. NPC2 facilitates bidirectional transfer of cholesterol between NPC1 and lipid bilayers, a step in cholesterol egress from lysosomes. Proc Natl Acad Sci U S A. 2008;105: 15287–15292. doi:10.1073/pnas.0807328105

17. Thelen AM, Zoncu R. Emerging Roles for the Lysosome in Lipid Metabolism. Trends in Cell Biology. Elsevier; 2017. pp. 833–850. doi:10.1016/j.tcb.2017.07.006

18. Alpy F, Rousseau A, Schwab Y, Legueux F, Stoll I, Wendling C, et al. STARD3 or STARD3NL and VAP form a novel molecular tether between late endosomes and the ER. J Cell Sci. 2013;126: 5500–5512. doi:10.1242/jcs.139295

19. Wilhelm LP, Wendling C, Védie B, Kobayashi T, Chenard M-P, Tomasetto C, et al. STARD 3 mediates endoplasmic reticulum-to-endosome cholesterol transport at membrane contact sites. EMBO J. 2017;36: 1412–1433. doi:10.15252/embj.201695917

20. Charman M, Kennedy BE, Osborne N, Karten B. MLN64 mediates egress of cholesterol from endosomes to mitochondria in the absence of functional Niemann-Pick Type C1 protein. J Lipid Res. 2010;51: 1023–1034. doi:10.1194/jlr.M002345

21. German JB, Smilowitz JT, Zivkovic AM. Lipoproteins: When size really matters. Curr Opin Colloid Interface Sci. 2006;11: 171. doi:10.1016/j.cocis.2005.11.006

22. Carroll RG, Zasłona Z, Galván-Peña S, Koppe EL, Sévin DC, Angiari S, et al. An unexpected link between fatty acid synthase and cholesterol synthesis in proinflammatory macrophage activation. J Biol Chem. 2018;293: 5509–5521. doi:10.1074/jbc.RA118.001921

23. Fukumitsu S, Villareal MO, Onaga S, Aida K, Han J, Isoda H. α-Linolenic acid suppresses cholesterol and triacylglycerol biosynthesis pathway by suppressing SREBP-2, SREBP-1a and −1c expression. Cytotechnology. Springer; 2013. pp. 899–907. doi:10.1007/s10616-012-9510-x

24. Saraswathi V, Hasty AH. The role of lipolysis in mediating the proinflammatory effects of very low density lipoproteins in mouse peritoneal macrophages. J Lipid Res. 2006;47: 1406–1415. doi:10.1194/jlr.M600159-JLR200

25. Lookene A, Nielsen MS, Gliemann J, Olivecrona G. Contribution of the carboxy-terminal domain of lipoprotein lipase to interaction with heparin and lipoproteins. Biochem Biophys Res Commun. 2000;271: 15–21. doi:10.1006/bbrc.2000.2530

26. den Hartigh LJ, Altman R, Norman JE, Rutledge JC. Postprandial VLDL lipolysis products increase monocyte adhesion and lipid droplet formation via activation of ERK2 and NFκB. Am J Physiol - Hear Circ Physiol. 2014;306. doi:10.1152/ajpheart.00137.2013

27. Aflaki E, Doddapattar P, Radović B, Povoden S, Kolb D, Vujić N, et al. C16 ceramide is crucial for triacylglycerol-induced apoptosis in macrophages. Cell Death Dis. 2012;3. doi:10.1038/cddis.2012.17

28. Oteng AB, Ruppert PMM, Boutens L, Dijk W, Van Dierendonck XAMH, Olivecrona G, et al. Characterization of ANGPTL4 function in macrophages and adipocytes using Angptl4-knockout and Angptl4-hypomorphic mice. J Lipid Res. 2019;60: 1741–1754. doi:10.1194/jlr.M094128

29. Milosavljevic D, Kontush A, Griglio S, Le Naour G, Thillet J, Chapman MJ. VLDL-induced triglyceride accumulation in human macrophages is mediated by modulation of LPL lipolytic activity in the absence of change in LPL mass. Biochim Biophys Acta - Mol Cell Biol Lipids. 2003;1631: 51–60. doi:10.1016/S1388-1981(02)00355-4

30. Vercauteren D, Vandenbroucke RE, Jones AT, Rejman J, Demeester J, De Smedt SC, et al. The use of inhibitors to study endocytic pathways of gene carriers: Optimization and pitfalls. Mol Ther. 2010;18: 561–569. doi:10.1038/mt.2009.281

31. Cui W, Sathyanarayan A, Lopresti M, Aghajan M, Chen C, Mashek DG. Lipophagy-derived fatty acids undergo extracellular efflux via lysosomal exocytosis. Autophagy. 2021;17: 690–705. doi:10.1080/15548627.2020.1728097

32. Martins IJ, Mortimer B, Miller J, Redgrave TG. Effects of particle size and number on the plasma clearance of chylomicrons and remnants. J Lipid Res. 1996;37. Available: www.jlr.org

33. Lindqvist P, Ostlund-Lindqvist AM, Witztum JL, Steinberg D, Little JA. The role of lipoprotein lipase in the metabolism of triglyceride-rich lipoproteins by macrophages. J Biol Chem. 1983;258: 9086–9092. doi:10.1016/s0021-9258(17)44634-5

34. Goldberg IJ. Lipoprotein lipase and lipolysis: Central roles in lipoprotein metabolism and atherogenesis. Journal of Lipid Research. 1996. pp. 693–707. Available: https://pubmed.ncbi.nlm.nih.gov/8732771/

35. Stein Y, Stein O. Lipoprotein lipase and atherosclerosis. Atherosclerosis. 2003;170: 1–9. doi:10.1016/S0021-9150(03)00014-5

36. Zheng C, Murdoch SJ, Brunzell JD, Sacks FM. Lipoprotein lipase bound to apolipoprotein B lipoproteins accelerates clearance of postprandial lipoproteins in humans. Arterioscler Thromb Vasc Biol. 2006;26: 891–896. doi:10.1161/01.ATV.0000203512.01007.3d

37. Borén J, Lookene A, Makoveichuk E, Xiang S, Gustafsson M, Liu H, et al. Binding of Low Density Lipoproteins to Lipoprotein Lipase Is Dependent on Lipids but Not on Apolipoprotein B. J Biol Chem. 2001;276: 26916–26922. doi:10.1074/jbc.M011090200

38. Hendriks WL, Van Der Boom H, Van Vark LC, Havekes LM. Lipoprotein lipase stimulates the binding and uptake of moderately oxidized low-density lipoprotein by J774 macrophages. Biochem J. 1996;314: 563–568. doi:10.1042/bj3140563

39. Ostlund-Lindqvist AM, Gustafson S, Lindqvist P, Witztum JL, Little JA. Uptake and degradation of human chylomicrons by macrophages in culture. Role of lipoprotein lipase. Arteriosclerosis. 1983;3: 433–440. doi:10.1161/01.atv.3.5.433

40. Takahashi M, Yagyu H, Tazoe F, Nagashima S, Ohshiro T, Okada K, et al. Macrophage lipoprotein lipase modulates the development of atherosclerosis but not adiposity. J Lipid Res. 2013;54: 1124–1134. doi:10.1194/jlr.M035568

41. Merkel M, Kako Y, Radner H, Cho IS, Ramasamy R, Brunzell JD, et al. Catalytically inactive lipoprotein lipase expression in muscle of transgenic mice increases very low density lipoprotein uptake: Direct evidence that lipoprotein lipase bridging occurs in vivo. Proc Natl Acad Sci U S A. 1998;95: 13841–13846. doi:10.1073/pnas.95.23.13841

42. Tuohetahuntila M, Molenaar MR, Spee B, Brouwers JF, Wubbolts R, Houweling M, et al. Lysosome-mediated degradation of a distinct pool of lipid droplets during hepatic stellate cell activation. J Biol Chem. 2017;292: 12436–12448. doi:10.1074/jbc.M117.778472

43. Bartelt A, Bruns OT, Reimer R, Hohenberg H, Ittrich H, Peldschus K, et al. Brown adipose tissue activity controls triglyceride clearance. Nat Med. 2011;17: 200–206. doi:10.1038/nm.2297

44. Heine M, Fischer AW, Schlein C, Jung C, Straub LG, Gottschling K, et al. Lipolysis Triggers a Systemic Insulin Response Essential for Efficient Energy Replenishment of Activated Brown Adipose Tissue in Mice. Cell Metab. 2018;28: 644–655.e4. doi:10.1016/j.cmet.2018.06.020

45. Schlein C, Fischer AW, Sass F, Worthmann A, Tödter K, Jaeckstein MY, et al. Endogenous Fatty Acid Synthesis Drives Brown Adipose Tissue Involution. Cell Rep. 2021;34: 108624. doi:10.1016/j.celrep.2020.108624

46. Fischer AW, Jaeckstein MY, Gottschling K, Heine M, Sass F, Mangels N, et al. Lysosomal lipoprotein processing in endothelial cells stimulates adipose tissue thermogenic adaptation. Cell Metab. 2021;33: 547–564.e7. doi:10.1016/j.cmet.2020.12.001

47. Maurer ME, Cooper JA. The adaptor protein Dab2 sorts LDL receptors into coated pits independently of AP-2 and ARH. J Cell Sci. 2006;119: 4235–4246. doi:10.1242/jcs.03217

48. Kibbey RG, Rizo J, Gierasch LM, Anderson RGW. The LDL receptor clustering motif interacts with the clathrin terminal domain in a reverse turn conformation. J Cell Biol. 1998;142: 59–67. doi:10.1083/jcb.142.1.59

49. Garuti R, Jones C, Li WP, Michaely P, Herz J, Gerard RD, et al. The modular adaptor protein autosomal recessive hypercholesterolemia (ARH) promotes low density lipoprotein receptor clustering into clathrin-coated pits. J Biol Chem. 2005;280: 40996–41004. doi:10.1074/jbc.M509394200

50. Ye ZJ, Go GW, Singh R, Liu W, Keramati AR, Mani A. LRP6 protein regulates low density lipoprotein (LDL) receptor-mediated LDL uptake. J Biol Chem. 2012;287: 1335–1344. doi:10.1074/jbc.M111.295287

51. Wei J, Fu ZY, Li PS, Miao HH, Li BL, Ma YT, et al. The clathrin adaptor proteins ARH, Dab2, and numb play distinct roles in Niemann-Pick C1-Like 1 versus low density lipoprotein receptor-mediated cholesterol uptake. J Biol Chem. 2014;289: 33689–33700. doi:10.1074/jbc.M114.593764

52. Jones NL, Willingham MC. Modified LDLs are internalized by macrophages in part via macropinocytosis. Anat Rec. 1999;255: 57–68. doi:10.1002/(SICI)1097-0185(19990501)255:1<57::AID-AR7>3.0.CO;2-Z

53. Rejman J, Oberle V, Zuhorn IS, Hoekstra D. Size-dependent internalization of particles via the pathways of clathrin- and caveolae-mediated endocytosis. Biochem J. 2004;377: 159–169. doi:10.1042/BJ20031253

54. Tabas I, Bornfeldt KE. Intracellular and Intercellular Aspects of Macrophage Immunometabolism in Atherosclerosis. Circ Res. 2020;126: 1209–1227. doi:10.1161/CIRCRESAHA.119.315939

55. Passeggio J, Liscum L. Flux of fatty acids through NPC1 lysosomes. J Biol Chem. 2005;280: 10333–10339. doi:10.1074/jbc.M413657200

56. Wilhelm LP, Tomasetto C, Alpy F. Touché! STARD3 and STARD3NL tether the ER to endosomes. Biochem Soc Trans. 2016;44: 493–498. doi:10.1042/BST20150269

57. Lichtenstein L, Mattijssen F, De Wit NJ, Georgiadi A, Hooiveld GJ, Van Der Meer R, et al. Angptl4 protects against severe proinflammatory effects of saturated fat by inhibiting fatty acid uptake into mesenteric lymph node macrophages. Cell Metab. 2010;12: 580–592. doi:10.1016/j.cmet.2010.11.002

58. Xie Z, Bailey A, Kuleshov M V., Clarke DJB, Evangelista JE, Jenkins SL, et al. Gene Set Knowledge Discovery with Enrichr. Curr Protoc. 2021;1: e90. doi:10.1002/cpz1.90

59. Chen EY, Tan CM, Kou Y, Duan Q, Wang Z, Meirelles G V., et al. Enrichr: Interactive and collaborative HTML5 gene list enrichment analysis tool. BMC Bioinformatics. 2013;14. doi:10.1186/1471-2105-14-128

60. Kuleshov M V., Jones MR, Rouillard AD, Fernandez NF, Duan Q, Wang Z, et al. Enrichr: a comprehensive gene set enrichment analysis web server 2016 update. Nucleic Acids Res. 2016;44: W90–W97. doi:10.1093/nar/gkw377

